# Catalytic cycling of human mitochondrial Lon protease

**DOI:** 10.1101/2021.07.28.454137

**Authors:** Inayathulla Mohammed, Kai A. Schmitz, Niko Schenck, Annika Topitsch, Timm Maier, Jan Pieter Abrahams

**Affiliations:** Biozentrum, University of Basel, Basel, Switzerland; Paul Scherrer Institute, Laboratory of Nanoscale Biology, Villigen, Switzerland

## Abstract

The mitochondrial Lon protease homolog (LonP1) hexamer controls mitochondrial health by digesting proteins from the mitochondrial matrix that are damaged or must be removed. Understanding how it is regulated requires characterizing its mechanism. Here, we show how human LonP1 functions, based on eight different conformational states that we determined by cryo-EM with a resolution locally extending to 3.6 Å for the best ordered states. LonP1 has a poorly ordered N-terminal part with apparent threefold symmetry, which apparently binds substrate protein and feeds it into its AAA+ unfoldase core. This translocates the extended substrate protein into a proteolytic cavity, in which we report an additional, previously unidentified Thr-type proteolytic center. Threefold rocking movements of the flexible N-terminal assembly likely assist thermal unfolding of the substrate protein. Our data suggest LonP1 may function as a sixfold cyclical Brownian ratchet controlled by ATP hydrolysis.

## Introduction

The proton potential gradient that drives mitochondrial ATP synthesis is established by electron transport chains that generate reactive oxygen species (ROS) as a side product (1). ROS compromises mitochondrial functionality by damaging proteins in the mitochondrial matrix. These are removed by the nuclear encoded unfoldase/protease LonP1, which is an important mitochondrial regulatory hub (2–4). LonP1 also binds mitochondrial DNA in a sequence specific manner (5), plays a vital role in mitochondrial DNA maintenance (6, 7), and regulates non-damaged proteins like ‘Transcription Factor A from Mitochondrial nucleoid’ (TFAM) (8). Even mild LonP1 (expression) deficiencies are linked to apoptosis and cell death (9) and several diseases are associated with dysfunctional LonP1, such as epilepsy, myopathy, paraplegia, cancer, and the CODAS (cerebral, ocular, dental, auricular and skeletal anomalies) syndrome (10, 11). Lon proteases are conserved over all kingdoms of life and belong to the HCLR clade of AAA+ unfoldases (12, 13).

A previous 21 Å resolution cryo-EM study of full-length human mutant LonP1 showed it has two globular parts. One part contains the N-terminal domains, and the other part contains the AAA+ and protease domains (14). Substructures of bacterial Lon protease from *Bacillus subtilis* (15), *Escherichia coli* (16) and *Meiothermus taiwanensis* (17) have been described in atomic detail. The substructure of the hexameric assembly of AAA+ and protease domains from *Yersinia pestis* indicated it is activated by inversion of the helical arrangement of its AAA+ domains (18). This and a similar study of human LonP1 (19) showed that protease activation closes a lateral gap between two subunits around an unfolded, extended chain of the substrate protein. As a result, tyrosine-valine motifs located on the loops of the AAA+- domain bind to the extended substrate protein strand in an axial tunnel. Previous studies indicated that the Lon proteases, including LonP1 contain a serine-lysine catalytic dyad (20–22), and histidine residues could potentially contribute to catalytic activity in *E. coli* LonP (23).

So far, no high-resolution structure of any full-length Lon protease or homolog was available. Hence it was unclear what the quaternary arrangement or role of the N-terminal domains is, or how the AAA+ domains unfold the protein substrate to feed it into the protease sites, or how unfolding is coupled to ATP hydrolysis and exchange. Here we present a comprehensive structure-function analysis of this human AAA+ protease, that was caught in action.

## Results

### Structure determination

We studied wild-type, *E. coli*-expressed human LonP1 by cryo-EM (see Materials and Methods for experimental details). Structures were determined in the presence of either an ATP/ADP mix, AMPPCP (adenosine-5’-[(β,γ)-methyleno]triphosphate, a nonhydrolyzable ATP analog), or ATPγS (adenosine-5′-[γ-thio]triphosphate, a slowly hydrolyzing ATP analog). In the latter case, we also included the substrate protein TFAM. These incubation conditions induced structurally distinct LonP1 species. Even on the same grid, multiple states were discerned. In total, we determined nine structures in eight conformational states (table 1, fig. S1-S3). Two of these states could only be determined at a resolution of around 12 Å. For the N-domains, only the overall domain shapes were apparent.

**Table 1.** Data collection and image processing of the respective datasets in different incubating conditions. In all cases, data were collected in movie-mode (40 frames/movie) at 300 keV on a Titan Krios TEM with a K2-camera in counting mode in a defocus range of - 1.0 to 2.5 μm, using a GIF energy filter and an energy dispersion of 20 eV.

|  | Dataset I | Dataset II | Dataset III |
| --- | --- | --- | --- |
| Included substrates | ADP + ATP | AMPPCP | ATPγS + TFAM |
| Nominal magnification | 165,000x | 130,000x | 130,000x |
| Camera mode | K2 (counting mode)<br>Super resolution<br>(8kx8k) | K2 (counting mode)<br>Super resolution<br>(8kx8k) | K2 (counting mode) |
| Pixel size | 0.831 Å | 1.058 Å | 1.058 Å |
| Exposure per movie | 8 s | 10 s | 10 s |
| # movies | 4100 | 3200 | 2520 |
| Total exposure | 80 e<sup>-</sup>/Å<sup>2</sup> | 80 e<sup>-</sup>/Å<sup>2</sup> | 60 e<sup>-</sup>/Å<sup>2</sup> |
| # particles (initial) | 280,000 | 320,000 | 198,000 |
| # particles per observed state | R-state: 82651<br>P1b-state: 31607<br>P1c-state: 47634<br>P2c-state: 30708 | R-state: 79850 | D1-state: 15000<br>D2-state: 11000<br>P1a-state: 62200<br>P2a-state: 26800 |
| Resolution masked maps (d99 / FSC at 0.143 of half maps) | R-state: 4.4 Å / 3.8 Å<br>P1b-state: 5.0 Å / 4.4 Å<br>P1c-state: 4.2 Å / 3.8 Å<br>P2c-state: 5.8 Å / 5.8 Å | R-state: 4.2 Å / 3.9 Å | D1-state: 12.6 Å / 14 Å<br>D2-state: 12.1 Å / 10 Å<br>P1a-state: 4.3 Å / 3.8 Å<br>P2a-state: 7.6 Å / 7.5 Å |

We improved resolution by focused refinement of the A- and protease domains and map-sharpening. However, focused refinement reduced the resolution of the N-domains. Separate focused refinement of the N-domains did not result in an improvement of their resolution. This indicated that the quaternary structure of the N-domains was more variable, even within the same state.

### Model building

For the A- and protease domains, well-resolved atomic models of homologous domains were used for initial atomic interpretation of the reconstructed scattering potential maps: 4ypl.pdb and 4fw9.pdb (the A- and protease domains of *M. taiwanensis* LonP). Subdomains of these structures were fitted as rigid bodies and mutated into the human mitochondrial LonP1 sequences. First the high resolution LonP1 domains were built. These were then used for interpreting more poorly ordered density. Missing sequences were built interactively. Where the local resolution was ambivalent, interactive model building was guided by density of other subunits or domains that showed clearer density. Torsion angles were restrained using non-crystallographic symmetry in Phenix (24) and significant deviations were investigated. Where appropriate, these were corrected interactively by considering all other subunits in all states. Great care was taken that the models of the different states were consistent with each other, down to main chain geometry. Real-space refinement included hydrogen atoms to improve geometry by better treating potential clashes.

The resolution of the A-domain nucleotide binding sites was insufficient to observe nucleotide phosphates as separate densities within the any of the scattering potential maps. To prevent refinement from unreasonably distorting the phosphate moieties into density, regardless of whether ATP or ADP was being fitted, we imposed four additional, strict distance restraints between main chain amides and phosphate oxides. These restraints were taken from a highly resolved adenosine nucleotide binding P-loop structure with good local sequence homology to LonP1: 2cbz.pdb, the human Multidrug Resistance Protein 1 Nucleotide Binding Domain 1 at 1.5 Å resolution. The following additional restraints were introduced in real-space refinement by Phenix: distance restraints of 2.64, 2.97, 3.33 and 2.66 Å between the main chain amides of G526, V527, G528 and T530 and phosphate oxygens O2β, O2β, O2α and O2α, respectively. In the case of ATP or ATPγS, additional distance restraints were introduced: 2.08 Å between the Mg^++^-ion and the Oγ1 oxygen of T530, and the phosphate oxygens Oγ1 and Oβ1; angular restraints secured orthogonal coordination of the Mg^++^-ion. These restraints ensured that all nucleotides were modelled consistently, with the same geometry as a 1.5 Å homologous structure.

Where the local resolution did not warrant atomic refinement, we identified the parts that behaved as rigid bodies, based on their domain topology and by scrutinizing superpositions with better resolved models from other, related states, and only refined their orientations. The hinge regions (e.g., fig. 2, S5, S6) connecting the rigid bodies were refined using torsion angle restraints derived from high resolution models, or equivalent regions from better resolved parts of the map. Where maps had a poor (local) resolution, this was reflected by high temperature values, which were refined per amino acid residue.

The globular parts of the N-domains (residues 123-353) had a poor local resolution of 8 to 12 Å in all observed states. Although this did not allow atomic interpretation, the overall quaternary arrangement of these N-domains and their orientation relative to the A- and protease domains could unequivocally be observed in all experimental maps (fig. S4B). These orientations differed between states. To characterize these quaternary rearrangements, homology models of the N-domains were fitted into their low-resolution density as rigid bodies. A humanized homology model of the *E. coli* N-domain (3ljc.pdb) that was calculated using SWISS-MODEL (25). The well-structured parts of the homology model had reasonable statistics (22.4 % sequence identity, a QMEAN of -2.53 and a GQME of 0.59; see Materials & Methods, table 2 and fig. S4B). Independently, and without imposing the *E. coli* N-domain template, I-TASSER (26) identified this structure as the closest equivalent and generated a very similar model with a C-score of -1.35 and a TM-score of 0.55. The model of the N-domain of human LonP1 that was generated independently of any homology models by AlphaFold (27; https://alphafold.ebi.ac.uk) also had the same fold (fig. S4C). All three models indicated residues 355-407 to form a long α-helix extending away from the globular domain and residues 1-123 and 221-272 to form unstructured loops. The structured cores of the I-TASSER and AlphaFold models deviated by 2.2 and 3.9 Å root mean squared difference (RMSD), respectively, from the model produced by SWISS-MODEL. The structured cores of all three models had the same overall shape, which fitted well within the low-resolution contours of all observed maps. We proceeded with the N-domain model that was produced by SWISS-MODEL.

**Table 2.** Refinement statistics. Each complete model contained 75×10^3^ atoms, of which 38×10^3^ were hydrogen. The refined models did not contain cis-Pro residues or twisted peptide planes. Model statistics of the well-ordered A- and protease domains are tabulated, *except for the D-states, for which we give the statistics of the full model, as all domains are approximately equally poorly ordered. The N-domain assemblies in the various states were significantly less well ordered and were therefore fitted as rigid bodies with close-to-ideal geometry. Statistics are shown for the N-terminal assembly with perfect threefold symmetry and optimized rotamers, prior to fitting in the maps of the various states and subsequent rigid body tweaking, which did not substantially alter the model statistics.

|  | R-state in<br>ATP & ADP | R-state in<br>AMPPCP | D1-state* | D2-state* | P1a-state | P1b-state | P1c-state | P2a-state | P2c-state | N-domains |
| --- | --- | --- | --- | --- | --- | --- | --- | --- | --- | --- |
| Co-factors |  |  |  |  |  |  |  |  |  |  |
| ATP |  |  |  |  |  | 4 | 5 | 5 | 5 |  |
| ATPγS | | | | | 5 | | | | | |
| ADP | 6 | 6 | 6 | 6 | 1 | 2 | 1 | 1 | 1 |  |
| Mg$^{++}$ | | | | | 5 | 4 | 5 | 5 | 5 | |
| Bonds |  |  |  |  |  |  |  |  |  |  |
| length (#>4σ) | 0.013 Å (4) | 0.015 Å (4) | 0.012 (0) | 0.012 Å (0) | 0.013 Å (1) | 0.011 Å (0) | 0.014 Å (2) | 0.011 Å (2) | 0.011 Å (0) | 0.004 Å (0) |
| angles (#>4σ) | 1.78° (23) | 1.68° (14) | 1.50° (36) | 1.50° (31) | 1.55° (5) | 1.60° (7) | 1.60° (6) | 1.79° (28) | 1.67° (25) | 0.81° (0) |
| MolProbity score | 1.97 | 2.10 | 2.19 | 2.21 | 1.91 | 1.88 | 1.82 | 1.95 | 1.87 | 1.74 |
| Clash score | 8.02 | 11.00 | 14.07 | 14.93 | 7.34 | 8.05 | 6.67 | 8.43 | 8.07 | 5.07 |
| Ramachandran |  |  |  |  |  |  |  |  |  |  |
| outliers | 0.03 % | 0.03 % | 0.06 % | 0.06 % | 0.03 % | 0.0 % | 0.03 % | 0.03 % | 0.00 % | 0.0 % |
| allowed | 9.55 % | 9.67 % | 9.35 % | 9.33 % | 8.64 % | 6.78 % | 7.03 % | 8.15 % | 6.62 % | 7.6 % |
| preferred | 90.42 % | 90.30 % | 90.59 % | 90.61 % | 91.32 % | 93.22 % | 92.94 % | 91.82 % | 93.38 % | 94.4 % |
| Rama-Z (RMSD) |  |  |  |  |  |  |  |  |  |  |
| whole (N=3222) | 2.10 (0.15) | 2.22 (0.15) | 2.53 (0.12) | 2.54 (0.12) | 2.08 (0.15) | 1.86 (0.15) | 1.77 (0.15) | 2.10 (0.15) | 2.00 (0.14) | 3.98 (0.17) |
| helix (N=1293) | 2.34 (0.10) | 2.21 (0.10) | 2.06 (0.09) | 2.07 (0.09) | 2.36 (0.10) | 1.89 (0.11) | 1.92 (0.11) | 2.22 (0.10) | 1.86 (0.10) | 1.87 (0.16) |
| Sheet (N=448) | 3.91 (0.20) | 3.98 (0.20) | 3.71 (0.18) | 3.71 (0.18) | 3.80 (0.22) | 3.78 (0.22) | 3.71 (0.23) | 3.89 (0.22) | 3.90 (0.22) | 3.44 (0.35) |
| Loop (N=1481) | 0.60 (0.19) | 0.43 (0.19) | 0.47 (0.015) | 0.48 (0.15) | 0.51 (0.18) | 0.47 (0.18) | 0.44 (0.18) | 0.36 (0.18) | 0.30 (0.18) | 3.21 (0.19) |
| Rotamer outliers | 0.04 % | 0.04 % | 0.02 % | 0.05 % | 0.00 % | 0.07 % | 0.11 % | 0.04 % | 0.00 % | 0.00 % |
| Cβ outliers | 0.00 % | 0.00 % | 0.00 % | 0.00 % | 0.00 % | 0.00 % | 0.00 % | 0.13 % | 0.07 % | 0.00 % |
| CaBLAM utlier | 2.84 % | 2.99 % | 3.30 % | 3.39 | 2.93 % | 2.84 % | 3.18 % | 2.99 % | 2.77 % | 3.19 % |
| ADP (B-factors)<br>min/max/mean |  |  |  |  |  |  |  |  |  |  |
| protein | 89/355/167 | 55/496/171 | 108/999/941 | 89/999/850 | 100/229/136 | 135/560/226 | 84/218/124 | 65/548/198 | 180/550/283 |  |
| ligand | 107/281/151 | 86/276/139 | 484/999/914 | 433/999/826 | 103/247/136 | 159/298/218 | 96/135/113 | 100/269/174 | 233/337/275 |  |
| FSC 0/0.143/0.5 | 3.3/3.8/3.9 Å | 3.7/3.8/4.0 Å | >12 Å | >12 Å | 3.5/3.6/4.0 Å | 3.7/4.0/4.4 Å | 3.3/3.6/4.0 Å | 6.0/6.5/7.9 Å | 4.1/4.3/6.0 Å |  |
| CC (mask) | 0.77 | 0.84 | 0.54 | 0.64 | 0.83 | 0.81 | 0.84 | 0.74 | 0.82 |  |
| CC (box) | 0.79 | 0.90 | 0.85 | 0.88 | 0.86 | 0.88 | 0.82 | 0.71 | 0.87 |  |
| CC (peaks) | 0.63 | 0.80 | 0.38 | 0.48 | 0.74 | 0.66 | 0.70 | 0.38 | 0.63 |  |
| CC (volume) | 0.77 | 0.83 | 0.44 | 0.56 | 0.83 | 0.81 | 0.83 | 0.74 | 0.82 |  |
| Mean CC ligands | 0.73 | 0.79 | 0.74 | 0.62 | 0.82 | 0.78 | 0.84 | 0.75 | 0.80 |  |

The α-helix (residues 355-407) that connected the globular part of the N-domain to the A-domain was straight for even numbered subunits. It could be fitted into the tube-like density that surrounded the N-passage (see fig. 1 and fig. S11). However, in the odd-numbered subunits, this α-helix had to be kinked at residue His391 to make the N-domain fit the observed contours. We assembled these N-domains into a hexamer with perfect threefold symmetry, which fitted the low-resolution maps. Since SWISS-MODEL had selected rotamers without considering the quaternary structure of the N-domains, we had to re-optimize the rotamers in order to minimize inter-domain clashes using Phenix. The orientations of the individual N-domains were separately refined as rigid bodies in each of the nine maps, resulting in small deviations from threefold symmetry. Finally, the loops connecting the N-domains to the A-domains (residues 407-412) were built into density. In all subsequent real-space refinements, each N-domain was treated as a single rigid body.

**Figure 1.**
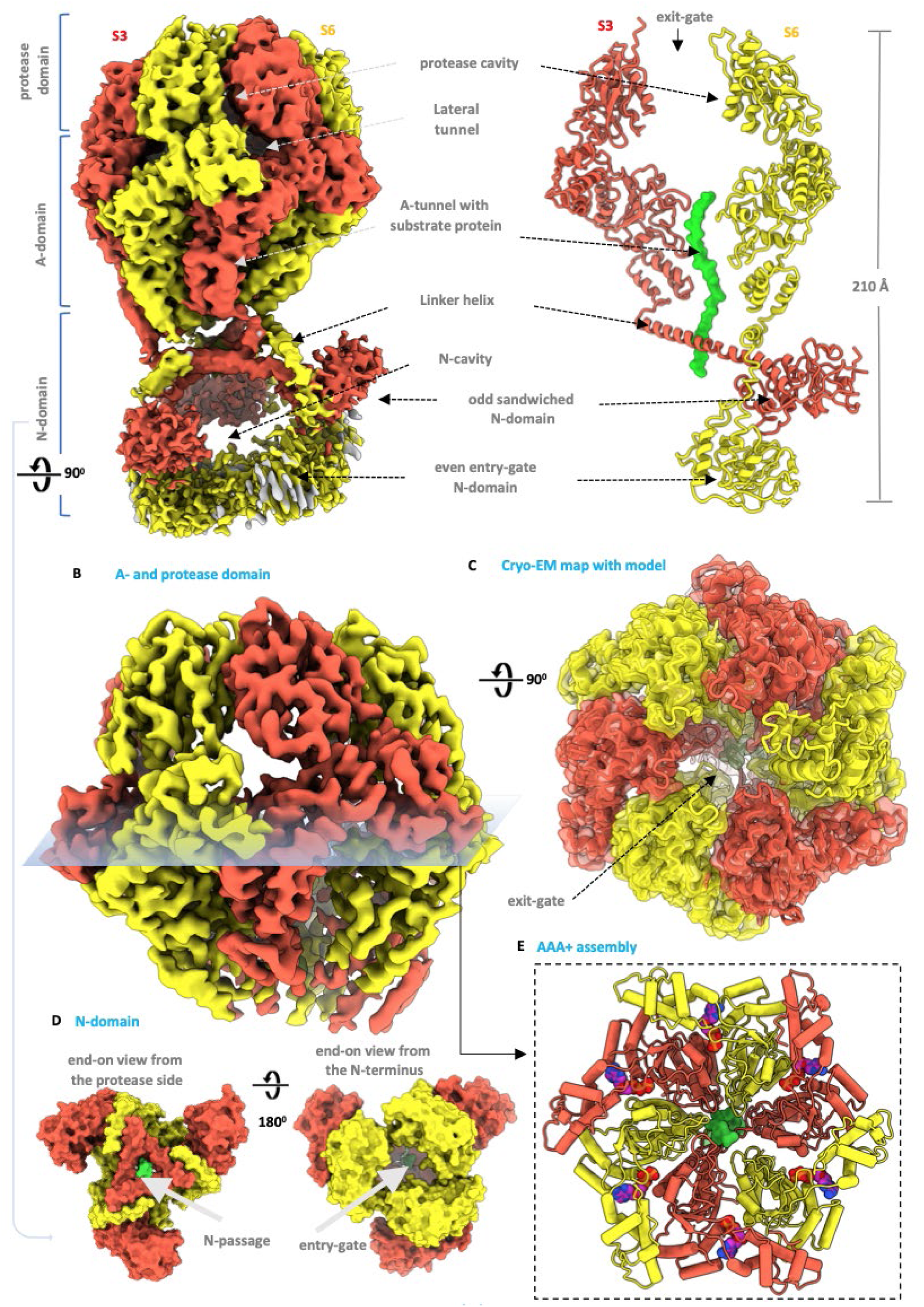
Structure of LonP1 in the P1a-state and nomenclature of its essential features. The odd- and even-numbered subunits are in red and yellow, respectively; the unfolded substrate protein strand is in green. (A) Lateral view of the scattering potential map (left), and of the atomic model with front and back subunits removed to show its internal structure and alternating arrangement of the odd- and even numbered N-domains (right). (B) Side view of the A- and protease domain scattering potential map, contoured at a higher level than in panel A, to illustrate the quality of the density, revealing the main chain fold, secondary structure, and side chain density of larger amino acids. (C) Top view of the scattering potential map, with the main chain indicated. (D) Top and bottom views of the trimeric N-domain assembly model (surface rendered). (E) Top view of the main chain trace of the A-domain assembly with α-helices and nucleotides indicated.

The refinement statistics of the real-space refined A- and protease domains of R- and P-states are listed separately from the statistics of the rigid body N-domain homology models (table 2). Some examples of secondary structural elements fitting into density are shown in fig. S4. All reported states differed significantly from each other in the well-ordered A-domain regions, with subunit shifts in the multi-Å range.

The D1- and D2-states had a poor overall map resolution of 14 and 10 Å, respectively (Table 1). We therefore took the R-state subunits and fitted these as rigid bodies into the D-state maps. We fitted the N-domains independently from the combined A- and protease domains, including bound nucleotides. We ensured the N-domains remained connected to the A-domains by modelling the loops 405 to 415 into density.

### Overall architecture

Our structures confirmed LonP1 has two globular parts (fig.1). Its poorly ordered bottom assembly of N-domains (residues 123-355) was similar in appearance to that of the 21 Å LonP1 structure reported earlier (14). The top had a local resolution up to 3.6 Å, allowing atomic model building of the ATP-binding A-domains (residues 414-749) and protease domains (residues 750-959). The nucleotide binding pockets were located at the subunit interfaces between adjacent A-domains, in the cleft between the α*/*β*−* and α-subdomains. The threefold axis of the N-domain assembly was tilted by up to 30° relative to the helical axis of the A-domains. A long linker-helix (residues 355-408), either straight or kinked, connected each N-domain to its corresponding A-domain.

The subunits (numbered 1 to 6) adopted cyclically alternating configurations. The linker-helix of each odd-numbered subunit was straight, and its corresponding N-domain was sandwiched between the N- and A-domains of the opposite even-numbered subunit. The N-domain assembly therefore was a trimer of stacked dimers. The linker-helix of each even-numbered subunit kinked round the linker-helix of the opposite odd subunit. The assembly of N-domains surrounded a 3×10^4^ Å^3^ N-cavity, accessible through three 40 Å lateral portals and an entry gate, formed by the bottom trimer of the even-subunit N-domains. The straight linker helices of the odd subunits formed a 10-15 Å wide N-passage, allowing access to an axial A-tunnel formed by the α*/*β-subdomains that is 8-10 Å wide. In the enzymatically active states that could be determined at sufficiently high resolution, we observed extended density in the A-tunnel, corresponding to a substrate protein that was being unfolded and digested. The A-tunnel led towards a 50 Å wide and 25 Å high protease cavity harboring the proteolytic sites, with a large exit gate at its apex and six smaller lateral tunnels. The A-domains were held together by the α-subdomains, which contained a conserved sensor-2 motif that facilitates ATP hydrolysis in related proteins (28, 29).

### Subunit conformations

Subunit conformations could differ significantly, mainly due to rigid body motions of their (sub-)domains. The A-domains were present in three different conformations. Based on their dominant nucleotide occupancy (see below), we refer to these conformations as the ATP-binding T-conformation, the ADP-binding D-conformation and unique S-conformation of the second seam subunit (see also fig. 2 and fig. S5, S6). Compared to the T-conformation, the α-subdomain of the D-conformation had tilted towards the helical axis by 9° to 13°. In the S-conformation, it had tilted downwards by a similar angle towards the α*/*β-subdomain. We quantified these angles by calculating the angular difference between the rotation operators that superimposed α*/*β-subdomains and the α-subdomains, respectively. The three-helical bundle of the α*/*β-subdomains showed small rigid body motions. Glycine-containing flexible loops attached the α- and α*/*β-subdomains to the protease and N-domains, respectively (fig. 2A). The orientation and packing of the protease domains and the poorly ordered stack of N-domains adapted to the conformation of the A-domains (movie 1).

**Figure 2.**
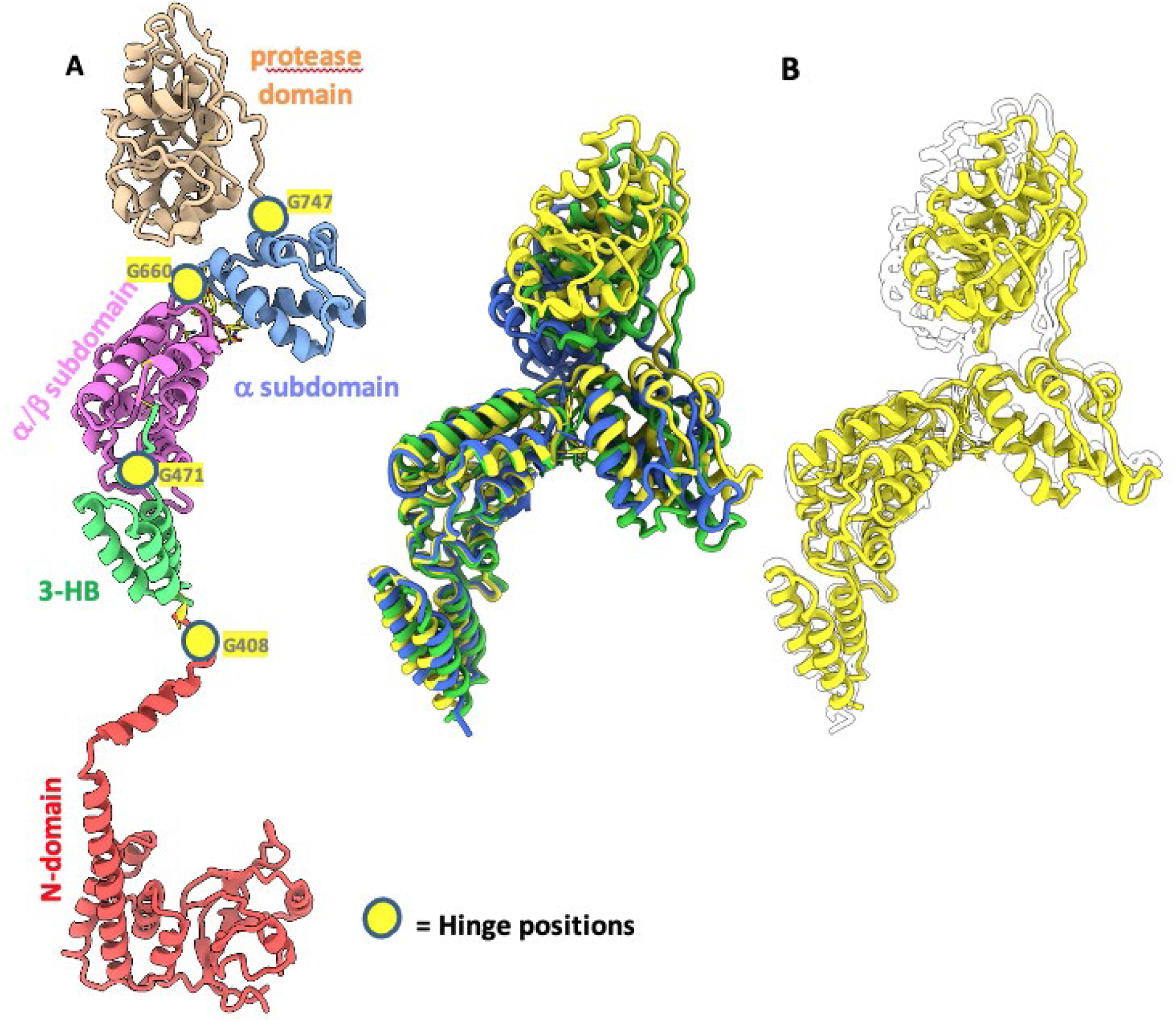
Hinge positions and the three major conformations of the A-domains. (A) Even subunit fold showing the individual subdomains in different colors, each linked with flexible glycine-containing loop, allowing rigid body movements. (B) Left panel: superimposition of α*/*β-subdomains from the P1a-state in the three major conformations encountered: the T-conformation (subunit 1; yellow), the D-conformation (subunit 5; blue) and the S-conformation (subunit 6; green). Right panel: superimposition of all A-domains from the P1a-state in the T-conformation (subunit 1 in yellow, subunits 2, 3 and 4 in white).

Compared to the D- and T-conformations, the S-conformation showed a small distortion of the central β-sheet of the α*/*β-subdomain. This sheet harbors two cysteines in parallel β-strands close to the P-loop (C520 on β1 and C637 on β4), that are conserved amongst mitochondrial proteases. In the D- and T-conformations, the distance between the Cβ-atoms of these cysteines was about 3.8 Å, which might allow disulfide linkage under oxidizing conditions. In the S-conformation, shearing of the central β-sheet increased this distance to about 5 Å, which would severely strain the geometry of a disulfide linkage (fig. 3A,B). Although disulfide formation has so far not been reported in the mitochondrial matrix, this observation may indicate a potential regulatory link between mitochondrial pathways that remove oxidizing species (30, 31), and LonP1 activity.

**Figure 3.**
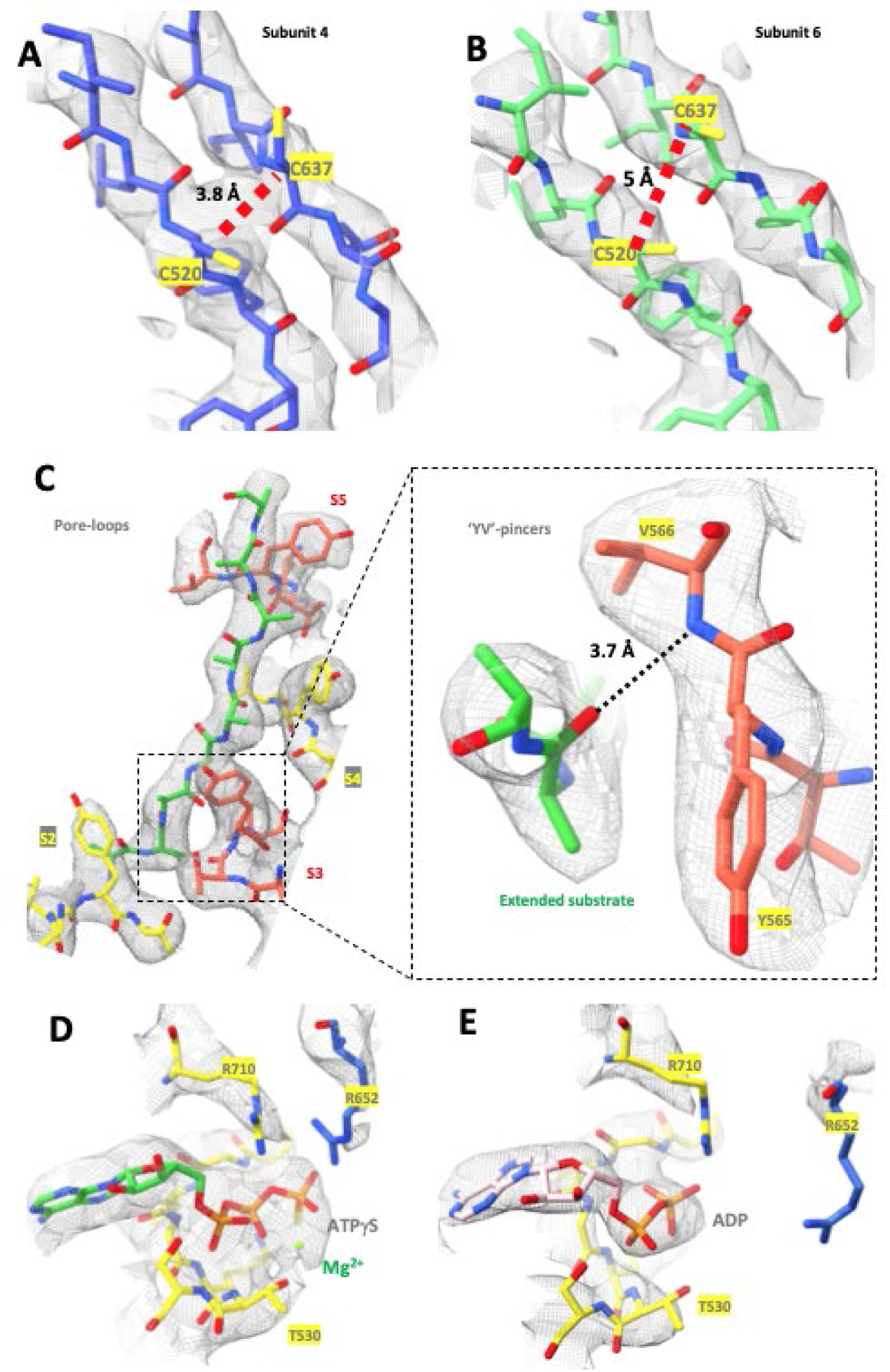
Map and model details. (A) Potential C520-C637 disulfide formation in the P1a-state in last helical subunit (S4) of the P1a-state. (B) In the second seam subunit, shearing of the β-sheet in which C520 and C637 are located, increases the distance between the Cβ*−*atoms of the cysteines to about 5 Å. (C) Binding of the extended substrate protein in the A-tunnel by YV-pincers may involve a hydrogen bond between the main chain oxygen of the substrate protein and the main chain amide between the Y- and V-residues of LonP1. (D, E) Nucleotide binding site and scattering potential map of subunit 4, showing ATPγS binding in the P1a-state (D) and ADP binding in the P1c-state (E).

### Enzymatic states

LonP1 occurred in eight different hexameric (sub)states, depending on the incubation conditions (see above). These states depended on the nucleotides that were added (AMPPCP, an ATP/ADP mixture, or ATPγS), but even within a single incubation condition, cryo-EM data processing separated multiple (sub)states. We distinguished a resting R-state, two infrequent docking D-states and five processing P-states with extended density in the A-tunnel. The A- and protease assemblies of the R- and P-states roughly corresponded to the LON^OFF^ and LON^ENZ^ states observed in a bacterial Lon protease (18).

The R-state was observed in the presence of ATP/ADP or AMPPCP (fig. 4 and table 1). Each A-domain adopted the D-conformation, which induced a left-handed pseudo-helical arrangement of the A- and protease domains with a twist and rise of about 55° and 5 Å per subunit. This caused a 30 Å axial displacement and a 23° wide gap, located towards the left of a subunit with a kinked linker helix, and which exposing an empty, 18 Å wide A-tunnel. Even when LonP1 was incubated in a buffer that exclusively contained AMPPCP, we only observed bound ADP, which suggests that LonP1 does not bind AMPPCP. The subunit that exposed its nucleotide binding site to the gap was less well-ordered than the other subunits. In the presence of AMPPCP, its main chains could still be traced, and its nucleotide density corresponded to ADP. However, in the presence of an ATP/ADP mixture, the contours of the α-subdomain and protease domain of the poorly ordered gap subunit were only visible at 1.8 and 2.8 RMSD, respectively, whereas for the other subunits, density of the main chain was visible at 6 RMSD. These poorly ordered domains were therefore docked as rigid bodies. We propose that in the R-state, this gap subunit may exchange ADP for ATP, but not for AMPPCP. This would result in multiple conformations for these domains in the presence of both ADP and ATP, explaining why under these conditions poor density for this subunit was observed.

**Figure 4.**
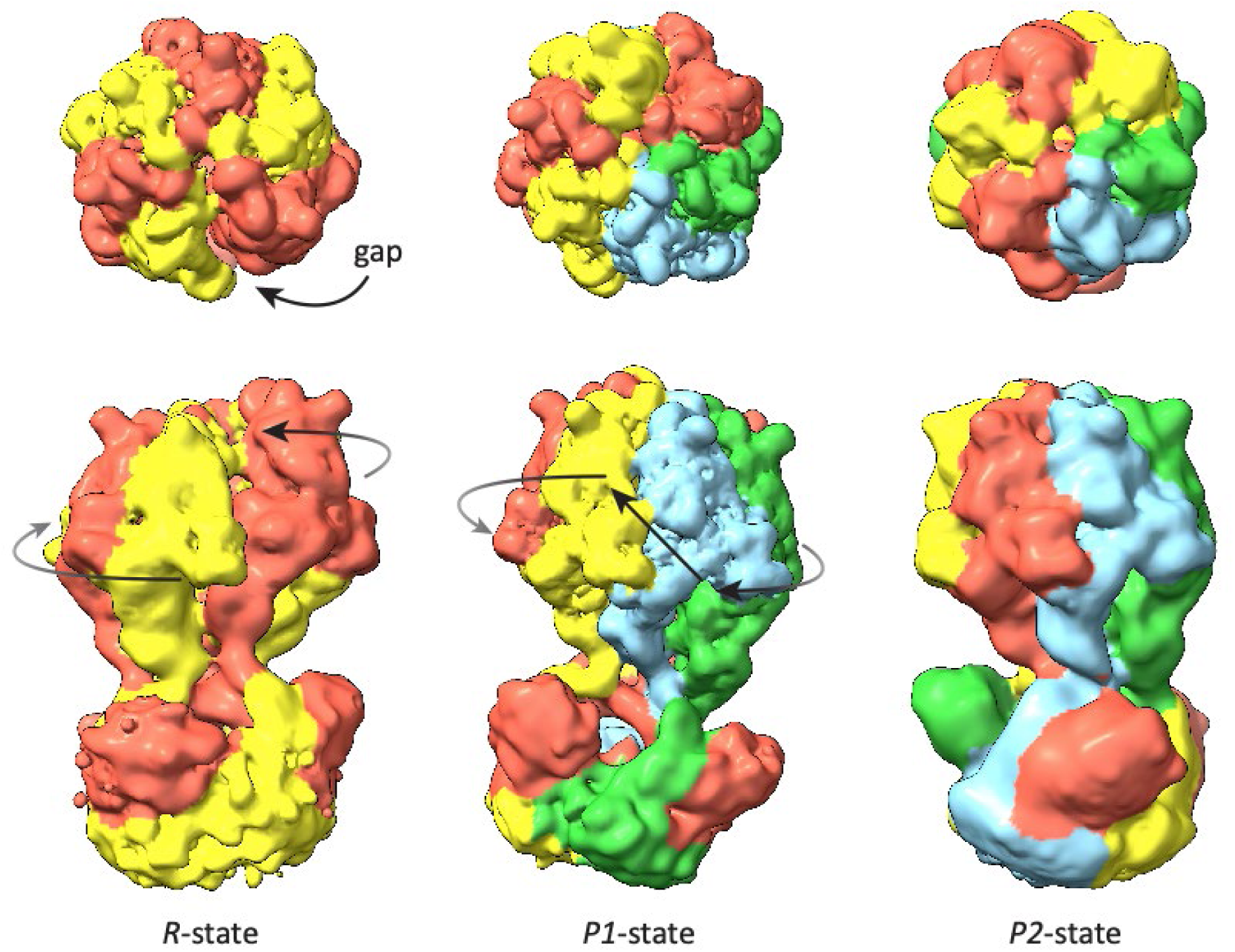
Locally denoised experimental maps of the R- and P-states of LonP1. The R-state has a lefthanded helical arrangement of the A- and protease-domains (indicated by curved arrows) and a gap to the left of an even-numbered subunit with a kinked linker helix (top left panel; even subunits in yellow, odd subunits in red). In the processing P-states the gap is closed, and four A-domains adopt a righthanded helical arrangement (indicated by curved arrows). Closing of the gap forces the two seam subunits (in blue and green) to compensate the righthanded helical rise by a left-handed arrangement (indicated by a straight arrow). In the P2-state, the N-terminal assembly appears to be rotated by 60° relative to the seam subunits. The maps were denoised by LAFTER (56).

The D1- and D2-states were observed after incubation with the slowly hydrolyzing ATPγS and the specific substrate protein TFAM. These states were uncommon and due to the low number of observed particles could only be resolved to 14 and 10 Å, respectively. In the D1-state, the R-state gap was not closed completely, and the A- and protease domains of the subunit left of the gap were particularly poorly ordered: their contours were only visible at 2.5 RMSD, whereas the contours of the other domains were clearly visible at 7.5 RMSD. In this respect, the D1-state resembled the R-state in the presence of ATP and ADP. In the D2-state, the gap had closed further, and the gap subunit was better ordered, with density at a contour of 4.5 RMSD, while the other subunits contoured at 7.5 RMSD. The D2-state showed density traversing the N-cavity (fig. 5), presumably corresponding to substrate protein. Although the resolution of the D-states did not allow identifying whether nucleotides were bound, we assumed this to be the case, because the highest scattering potential (>12.5 RMSD) was located at the nucleotide binding sites. We modelled all bound nucleotides as ADP, but some were most likely ATPγS since the D-states were only observed in the presence of this nucleotide analog.

**Figure 5.**
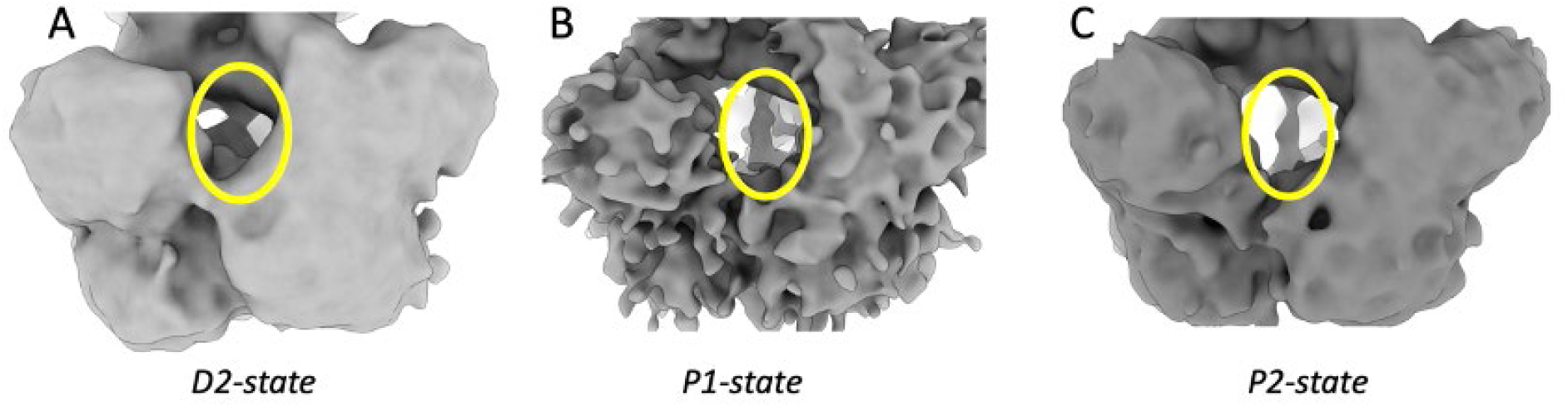
Extended substrate protein density in the N-cavity. The D2-state map (A) and P1- and P2-state maps that were not subjected to focused refinement on the A- and protease domains of the P1- and P2-states (B and C, respectively) show extended extra density within the N-cavity (highlighted in yellow), that rises from the entry gate to the N-passage towards better-ordered density within the A-tunnel. No non-crystallographic symmetry averaging of the N-domain assembly was imposed.

In the P-states, the R-state gap had closed. We observed two major P-states, each with sub-states. The P1-states were more abundant than the P2-states and differed by a major re-orientation of the AAA+ region relative to the N-terminal domain assembly. When the A- and protease densities of the P1-state were optimally superimposed on the corresponding densities of the P2-states, the N-domain densities of the two states were rotated by 60° with respect to each other (fig. 4). In the well-ordered P-states, extended density within the A-tunnel could be modeled as a poly-Ala chain with a slight right-handed helical twist (fig. 3C). The direction of the extended substrate protein chain was unclear.

Maps that were not subjected to focused refinement on the A- and protease domains had a lower resolution, as they were not dominated by the alignment of the well-ordered A- and P-domains. However, presumably because in these maps the more poorly ordered N-domains contributed stronger to the averaged scattering potential, the intermediate maps showed more clear density for the N-domain assembly. Like in the D2-state map, extended substrate protein density was visible in the unfocussed P-state maps, that emerged from the N-domain entry tunnel and traversed the N-cavity and the N-passage into the A-tunnel (fig. 5). We therefore propose that the substrate protein traverses the N-cavity.

In the P-states, four adjacent A-domains were present in the T-conformation. These subunits adopted a right-handed helical arrangement of the A-domains with an average helical twist and rise per subunit of about 60° and 7 Å, respectively (fig. 4 and fig. S5). To distinguish these four subunits from the seam subunits, we refer to them as the helical subunits. The A-domains of the two seam subunits were translated downwards to compensate for the rise of the helical subunits. The first seam subunit adopted the D-conformation that is also present in the R-state, and the second seam subunit adopted the S-conformation. In the P1-states, subunits 5 and 6 were the seam subunits, while in the P2-states, subunits 4 and 5 were the seam subunits. In the P2-state, the N-domain assembly therefore appeared to be rotated by 60° relative to the helical subunits, compared to the P1-state (fig. 4).

Based on significant domain movements, we distinguished substates P1a, P1b and P1c within the P1-state and substates P2a and P2c within the P2-state (we did not observe a P2b-state). The P1a- and P2a-states were observed after incubation with ATPγS and TFAM, while ATP/ADP induced the P1b-, P1c- and P2c-states. A comparison between the P-substates indicated that, to a large extent, the A-domains of the four helical subunits behaved as a single rigid group, with the seam subunit A-domains roughly acting as a second rigid group. Compared to the P1a-state, the seam subunits were displaced roughly downwards by about 3 and 6 Å in the P1b- and P1c-state, respectively. Similarly, the seam subunits had descended by about 6 Å in going from the P2a- to the P2c-state (fig. S6).

### Nucleotide binding

By introducing distance and angular restraints, we ensured all nucleotide phosphate moieties bound the P-loops as observed in high-resolution structures. We assigned the nucleotide based on the scattering potential at the γ-phosphate location. It was identified as ATP if a γ-phosphate was inside a contour corresponding to 2/3 of the observed scattering potential of either the Pα- or Pβ- phosphorus atom of the bound nucleotide (whichever was lower). In most cases this gave a reasonable contour for the di- or triphosphate moiety of the nucleotide, but at lower resolution this criterion was not always unambiguous (fig. S7). The proximity of local positive charges to the position of the γ-phosphate, especially of the trans-acting Arg652, correlated well with identified nucleotides. We exclusively observed ADP in the R-states. We do not exclude the possibility of a mixed ADP/ATP occupancy for some of the binding sites of the P-states. Nucleotide occupancies and subunit conformations are summarized in table 3.

**Table 3.** Nucleotide occupancies and conformational states of LonP1 subunits. The D-, T- and S-subunit conformations are highlighted in blue, yellow and green, respectively, and the identified nucleotide of each subunit is tabulated as a function of observed catalytic state (see fig. S6 for nucleotide scattering potentials of the P-states). All subunit conformations could be identified with certainty based on rigid body fitting and/or domain superimposition, except for subunit 4 in the R-state in the presence of ADP and ATP. The resolution of the P2a-state was insufficient for identifying the nucleotide occupancy and was assigned based on high structural similarity with the P1a-state after a 60° rotation. For subunit 4 of the P1a-state, the map did not exclude mixed occupancy. The conformations and nucleotide occupancies of the D-states did not allow unequivocal assignment of subunit conformations.

|  | S1 | S2 | S3 | S4 | S5 | S6 |
| --- | --- | --- | --- | --- | --- | --- |
| R-state in AMPPCP | ADP | ADP | ADP | ADP | ADP | ADP |
| R-state in AD/TP | ADP | ADP | ADP | ? | ADP | ADP |
| P1a-state | ATPγS | ATPγS | ATPγS | ATPγS | ADP | ATPγS |
| P1b-state | ATP | ATP | ATP | AD(T)P | ADP | ATP |
| P1c-state | ATP | ATP | ATP | ADP | ATP | ATP |
| P2a-state | ATPγS? | ATPγS? | ATPγS? | ADP? | ATPγS? | ATPγS? |
| P2c-state | ATP | ATP | ADP | ATP | ATP | ATP |

In the P-states, the additional scattering potential of a Mg^++^-ion coordinating the Oγ-atom of T530, the β- and the γ-phosphate when ATP was bound, contributed to differentiating between ADP and ATP. Although Mg^++^ is relatively light, its positive charge substantially increases its scattering potential. In the P1a-state, differentiating between ADP and ATPγS was further facilitated by the increased scattering potential of sulfur of the γ-thiophosphate moiety. In the P1-states, the first three helical subunits bound ATP (or ATPγS). The fourth helical subunit bound ATPγS in the P1a-state, in the P1b-state ADP fitted better than ATP, and the P1c-state ADP was bound. Subunit 5 bound ADP in the P1a- and P1b-states but appeared to bind ATP in the P1c-state. Subunit 6 appeared to bind ATP or ATPγS in all P1-states (fig. S7).

The P2-states were less common and had a lower resolution compared to the P1-states. The P2a-state required fitting subunit domains of the P1a-state as rigid bodies and did not allow unequivocally identifying its nucleotides. Because of the high similarity of the orientations of the A- and protease domains observed in the P1a- and P2a-states, we assumed that the nucleotide occupancy of these states also corresponded. The resolution of the P2c-state was just about enough to recognize that the nucleotide biding sites were occupied. In analogy to the P1c-state, density allowed us to place ATP in all binding sites of the P2c-state, except for subunit 3, which might have bound ADP (fig. S7).

In all P-states, the seam subunits had open sites potentially allowing ADP/ATP exchange. The ATP γ-phosphate of the first three helical subunits interacted with the trans-acting R652-finger of the next subunit (fig. S8), a motif also found in other AAA+ proteins. In the P1a- and P2a-states, the ATP γ-phosphate of the fourth helical subunit also interacted with this trans-acting R652-finger of the first seam subunit, whilst also the glutamate D612/E614-loop of the first seam subunit had approached (fig. S8). In other m-AAA proteases this activates ATP hydrolysis (32).

### Binding of substrate protein

In the high-resolution P-states, substrate protein density within the A-tunnel was observed in the presence or absence of the specific substrate protein TFAM. Although the main chain density of the substrate protein was reasonably well-ordered in the A-tunnel, there was no visible side chain density, and the resolution was insufficient to unequivocally determine the direction of the substrate protein chain. We investigated the peptides that were produced by active LonP1 by mass spectrometry. When we excluded TFAM, we found that LonP1 peptides were being produced. If we included another substrate protein in the reaction, their digested peptides were produced. For more details, see fig. 7C and supplementary materials (table S1B, table S2, fig. S10).

In all T-conformations of the P1-states, the conserved pore-loop1 residues Y565 and V566 of the helical subunits formed YV-pincers, that pointed into the A-tunnel along regularly spaced right-handed helical intervals and gripped the main chain of the stretched substrate protein along a length of 10 residues (fig. 6). The density and stereochemical considerations suggest a hydrogen bond between a main chain oxygen of the substrate protein strand and the main chain amide between the Y- and V-residues of the pincer may be involved in gripping the substrate protein main chain in a sequence independent manner (fig. 3C). In the P1a-state, the YV-pincer of the first seam subunit continued the helical rise and also gripped the substrate protein main chain. In the P1b- and P1c-states, the YV-pincer of the first seam subunit had released and together with the YV-pincer of the second seam subunit, had moved downwards along with the A-domains. The more poorly resolved P2-states showed less density for the substrate protein in the final maps that resulted from focused refinements. By analogy to the P1-states, we could partially trace the substrate protein through the A-tunnel in the P2c-state, where density for the substrate protein interacting with the four YV-pincers was visible at a contour level of 5 RMSD. The low resolution of the P2a-state prevented observing the substrate protein bound in the A-tunnel. However, also supported by the maps prior to focused refinement (cf. fig. 5), we assume the P2a- and P2c-states would interact in a similar fashion with the substrate protein as the P1a- and P1c-states.

**Figure 6.**
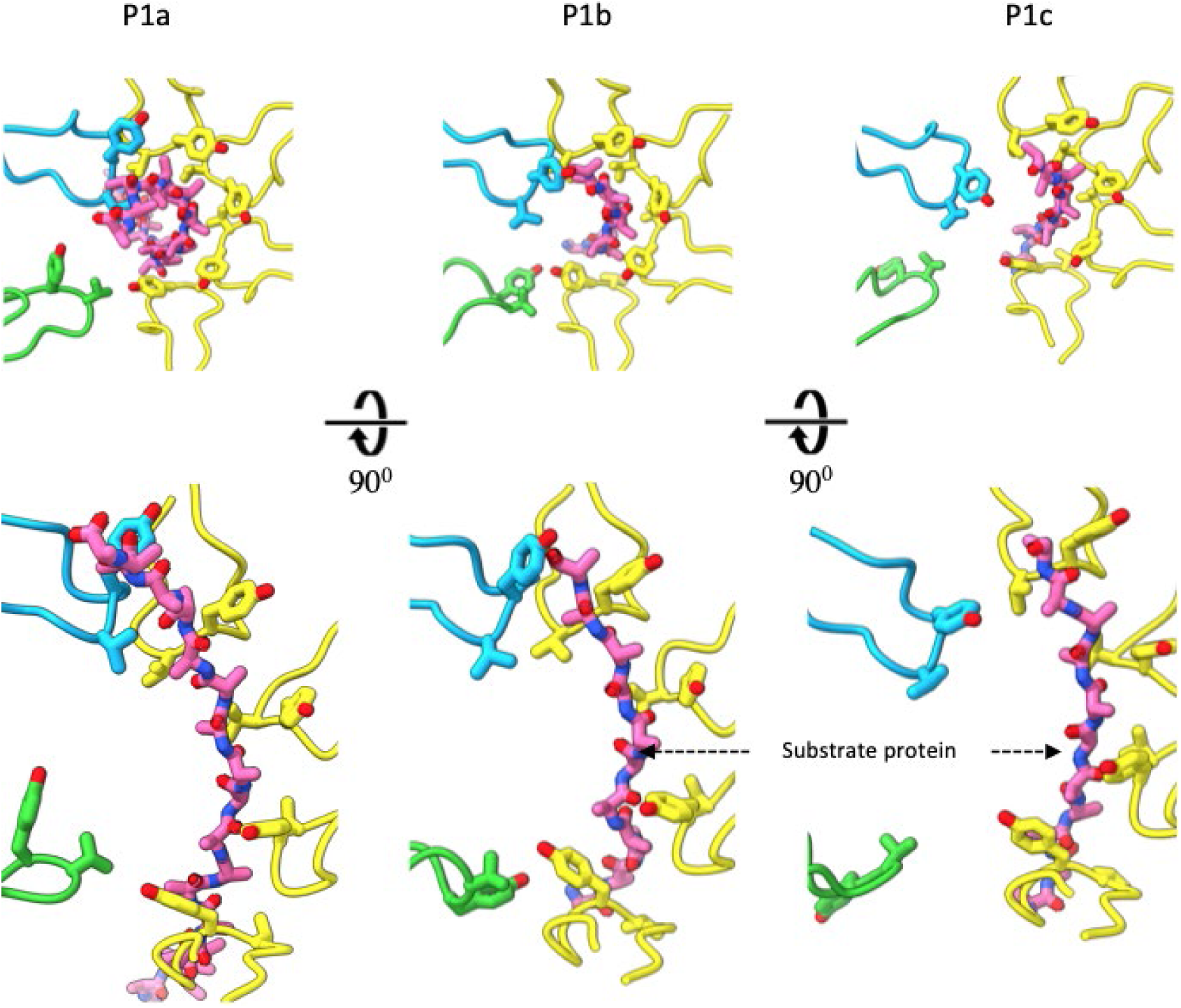
Interactions between the pore-loop1 YV-pincers and the extended substrate protein strand in the A-tunnel. In all P1-states, the YV-pincers of the helical subunits (in yellow) grab the mainchain of the substrate protein. In the P1a-state, the YV-pincer of the first seam subunit (in blue) grabs the top of the extended substrate protein strand below the protease cavity. In the P1b-state, this YV-pincer had released and the pore-loops of the seam subunits (in blue and green) had descended by about 3 Å. In the P1c-state, the pore-loops of the seam subunits had descended by another 3 Å, approximately. In the P2a- and P2c-states, very similar interactions are found.

### Enzymatic activity

Since ATP hydrolysis is activated in trans in AAA+ proteases, we used an adenosine triphosphate hydrolase (ATPase) assay as an additional control that our LonP1 construct formed functional hexameric complexes. We established that the level of ATPase activity of LonP1 was independent of the type of substrate protein, by assaying both the specific substrate protein TFAM and non-specific fluorescein isothiocyanate (FITC)-labeled casein. We confirmed that proteolysis requires ATPase activity by generating K529R and E591Q LonP1 mutants, replacing fully conserved residues of the Walker-A and -B motifs, respectively (33). These mutations indeed killed both ATPase and proteolytic activity, as determined by a fluorescence assays (fig. 7C, table S1A).

**Figure 7.**
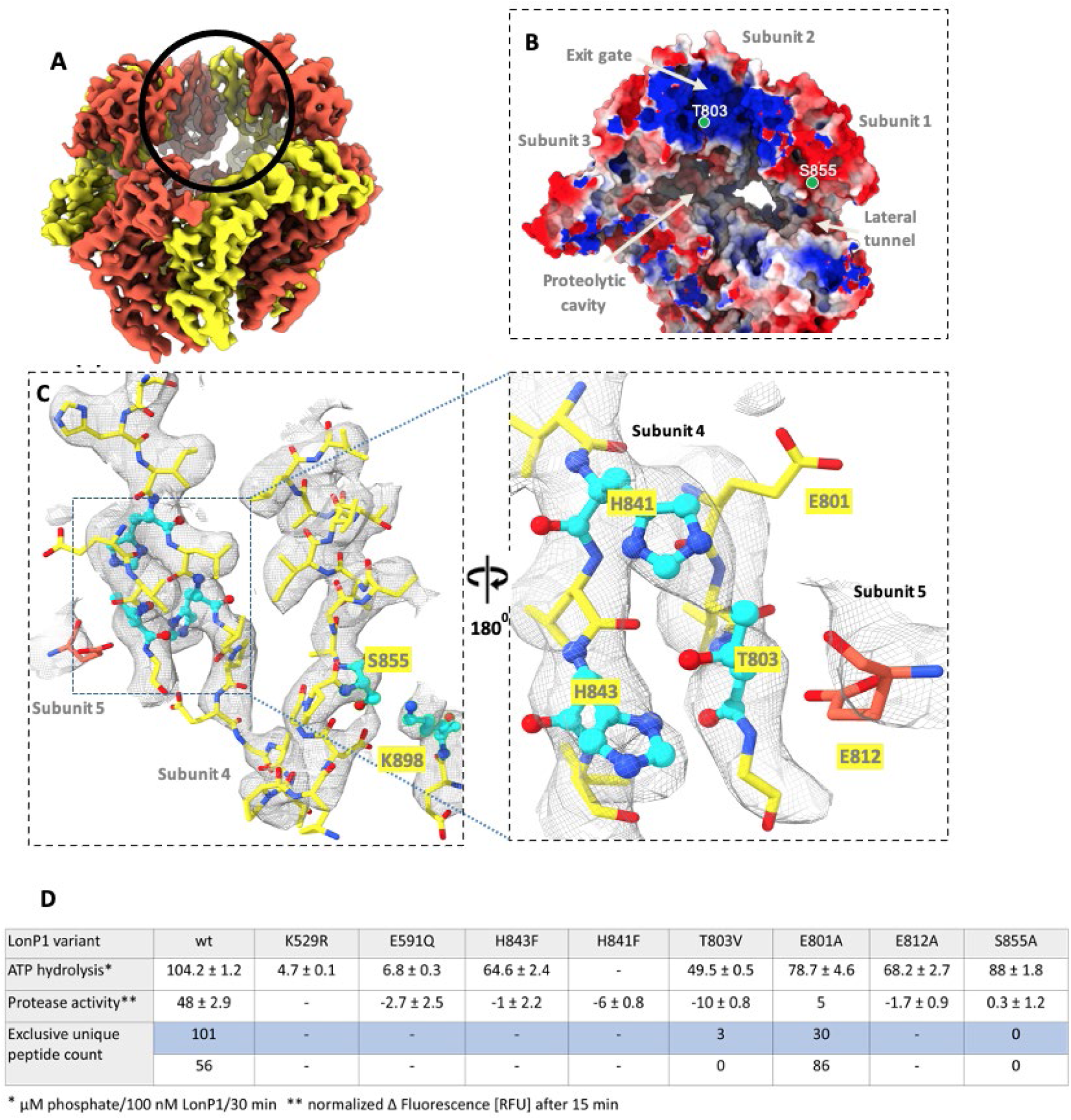
Structural arrangement of amino acids surrounding T803 that potentially participate in proteolysis. (A) Location of the conserved protease sites in the protease cavity is indicated by a black circle (for the sequence alignment, see fig. S9). The scattering potential map is colored with odd subunits in red and even subunits in yellow. (B) The proteolytic residue S855 is located within a recess of the lateral tunnel and the second, proposed proteolytic residue T803 is located at the subunit interface lining the apical exit gate. The positions of the S855 and T803 of subunits 1 and 2, respectively, are indicated with green spheres. The molecular surface is colored according to its surface potential (blue is positive, red is negative). (C) Close-up of the two proteolytic sites with a further close-up of the T803 proteolytic site, including H841 and E801. The S855-K898 catalytic dyad and the potential T803, H841 and H843 residues are depicted in ball-and-stick representation (cyan). (D) ATP hydrolysis and protease activity for LonP1 variants. FITC-casein was added to the ATP hydrolysis experiment. The peptide count of the mass spectrometry experiment is given (blue background: LonP1 peptides, white background: β-casein peptides).

We confirmed the S855/K898 catalytic dyad by generating the S855A mutation, which did not show any proteolytic activity, yet had an ATPase activity that was very close to that of wild-type LonP1. These six catalytic dyads were located within a small recess of the lateral tunnels, which had a surface charge that was mostly negative (fig. 7B).

Protease activity of bacterial Lon protease is activated by structural rearrangements around the catalytic S855/K898 dyad that are induced by the transition from the R-state to the P-state (18, 19). However, in our LonP1 structures, all individual R- and P-state proteolytic domains superimposed very well with RMS deviations of Cα-positions in the sub-Å range, and we did not observe full occlusion of the reactive S855 by other structural elements in any of the states, including D852, which has been identified as a regulatory element in bacterial Lon protease (21).

Based on a multiple sequence alignment (fig. S9) and after scrutinizing the LonP1 structure in all its states, we identified several other conserved residues within the protease domain that might be important for proteolytic activity. We considered T803, which is exposed in the proteolytic cavity, to be of special interest (fig. 7). This conserved residue is unlikely to be required for structural integrity because it does not appear to be involved in a hydrogen bonding network that maintains the tertiary or quaternary structure of the protease domains. However, its sidechain hydroxyl group can be rotated to within hydrogen-bonding distance of the imidazole groups of either H841 or H843, potentially forming a catalytic center. Furthermore, both H841 and H843 have an abundance of potentially activating acidic residues in their close environment and T803, H841 and E801 can form a catalytic triad. In many proteolytic triads, either serine or cysteine attacks the scissile peptide bond, but there are also examples of catalytic threonine (34–36). Finally, R815 is well positioned for moving its guanidinium moiety towards T803 and either act as the oxyanion hole or interact with T803’s hydroxyl group like in the S855-K898 catalytic dyad. All these residues are fully conserved amongst all mitochondrial proteases – with a unique exception for H843, which is a cysteine in platypus (*Ornithorhynchus anatinus*). The six potentially catalytic T803 residues were located on the surface of the large apical exit gate. The surface potential of the exit gate is positively charged, which may assist abstracting a proton from the γ-O of T830, which may enhance its proteolytic activity (fig. 7B).

The relevance of T803 for proteolysis was confirmed by the T803V mutant. The mutant T803V assembled into intact, ATP hydrolyzing hexamers – albeit at about 50 % of wild-type ATPase levels. However, changing the sidechain γ-hydroxyl group of T803 to a methyl group was sufficient to kill all proteolytic activity (fig. 7C). We further tested whether T803 might be the reactive residue of one or more catalytic triads, by mutating the potential bases H841 and H843 to phenylalanine, whilst retaining the wild-type T803. Also, these mutants hydrolyzed ATP but were proteolytically inactive, suggesting both residues were required and either might function as the base abstracting a proton from the T803 γ-OH moiety. We subsequently investigated which residues could potentially activate H841 or H843 by generating the E812A and E801A mutants, whilst retaining all other residues to wild type. The E812A mutant no longer showed proteolytic activity in fluorescence assays, whilst the E801A showed residual protease activity (fig. S10). Both mutants retained ATPase activity at about 70-75 % relative to the wild-type LonP1. A highly sensitive mass spectrometry assay confirmed that E801A was still capable of generating peptides in an ATP-dependent manner (fig. 7C, table S1B, table S2 and fig. S9). We conclude that LonP1 likely has a second proteolytic site, with T803 as the catalytic residue and with H841 or H843 as proton acceptors, which can be activated by conserved surrounding glutamates.

With these mutational studies, together with the possibility of forming a catalytic triad, T803 passes the same level of evidence as the S855/K898 catalytic dyad. It is not clear why LonP1 would have two different types of catalytic site. Our mutational studies showed that abolishing either of the two sites, whilst retaining the other, killed protease activity. The sites are too far apart both in *trans* and in *cis*, to consider them as a single active site (fig. 7B). It is not clear why both sites are required for proteolysis. It may be the case that the two sites have different specificities and need to act together to cleave the substrate protein into sufficiently small fragments to diffuse out of the protease cavity through the exit gate and prevent the protease cavity from clogging up. Or both sites have different functionalities, specific to either LonP1’s initiation or processing stages. Or killing one site might allow the substrate protein to escape the other site by diffusing either out of one of the lateral tunnels, or out of the apical exit gate.

## Discussion

Based on the high-resolution structures of the A- and protease domains, we propose a rotational catalytic cycle that accounts for all the observed states and even indicates why the N-domain assembly is more poorly resolved.

As a likely first step in LonP1 activation, the substrate protein binds the N-domain assembly, presumably in the R-state. As the N-domain density is poorly ordered, any speculation on the interactions between N-domains and the substrate protein are currently unwarranted and care should be taken that the N-domain homology model is not over-interpreted. We propose that either the C- or N-terminal sequence of the substrate protein is guided through the entry gate, N-cavity, and N-passage towards the A-tunnel. Rigid body fitting of R-state subunits into the D1-state map suggested that the YV-pincer of the left-hand, bottom R-state gap-subunit is translated upwards relative to the preceding subunit, whilst the R-state gap closes. This would require a trans-interaction of the gap-subunit R625-finger with ATP bound by its preceding, left-hand neighbor. Similar rigid body fitting into the D2-state map suggested that the next left-hand subunit is translated upwards by a similar mechanism (fig. S11). Subsequent conformational rearrangements that are required for reaching a P-state would involve translating the next two subunits downwards and grab the substrate protein. All these rearrangements are probably fast and dependent on ATP hydrolysis, since the D1- and D2-states were only observed in the presence of ATPγS.

The structures suggest that cyclic interconversions between P1- and P2-states drive substrate protein translocation by two amino acids per interconversion. This process requires the topmost YV-pincer to release the substrate protein strand, move downwards and then reattach at the bottom of the A-tunnel in a hand-over-hand mechanism that was proposed for bacterial Lon protease (18) and VAT (37) (a *T. acidophylum* AAA+ unfoldase). The five different P-states we present here, indicate an ATPase-dependent allosteric mechanism with cyclic threefold interconversions of states and sixfold interconversions of A-domain conformations (fig. 8, movies M1 and M2). Below, we discuss how the cyclic interconversion of P1- and P2-states may be regulated by ATP exchange and hydrolysis.

**Figure 8.**
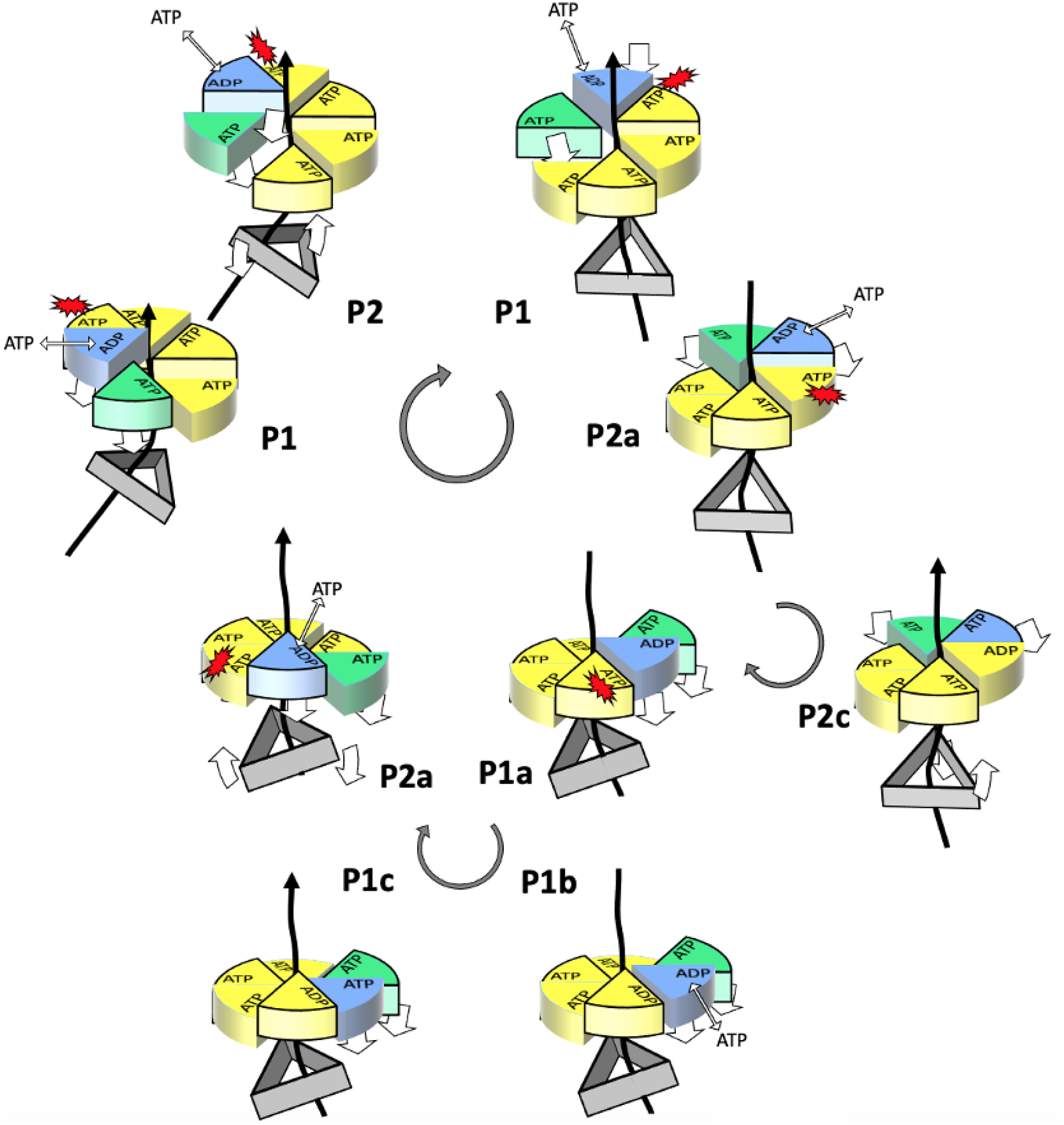
Diagram of the LonP1’s catalytic cycle. The A-domains and their helical arrangement are represented by cylindrical wedges with the bound nucleotide as indicated; the even-numbered subunits with a straight linker helix are shaded. A-domains in the D-conformation, the T-conformation and S-conformation are in blue, yellow and green, respectively. The tilt of the N-domain assembly is indicated by a grey triangular prism. The stretched-out substrate protein in the A-tunnel is represented by a solid black line. The upwards translocation of the substrate protein in the P1c-to-P2 and the P2-to-P1a transitions is indicated with a solid arrow. Domain movements are indicated with white arrows. The central cycle indicates the cyclical transitions between the P1- and P2-states. The peripheral bottom and left cycles indicate the transitions within the P1- and the P2-states, respectively.

Transition of the P1a- to the P1b-state requires the YV-pincer of subunit 5 (the first seam subunit) to release the substrate protein at the top of the A-tunnel. This transition is induced by ATP hydrolysis at subunit 4. ATP hydrolysis at subunit 4 is likely facilitated by the trans-acting D612/E615-loop of subunit 5 that in the P1a-state has moved towards subunit 4’s γ-phosphate (fig. S7). ATP hydrolysis releases the R652-finger of subunit 5 from the γ-phosphate moiety of nucleotide at subunit 4, allowing the downward motion of both seam subunits and concomitant release of the protein substrate by the YV-pincer of subunit 5. We propose ATP hydrolysis to be the rate limiting step in the P1a-to-P1b transition, since the P1a- (and P2a-states) could only be observed upon incubation with ATPγS, which hydrolyses 200 times more slowly than ATP.

Subsequent P1b-to-P1c transition results in a further downward movement of both seam subunits. We propose that this transition is facilitated by ADP/ATP exchange at subunit 5, because this subunit shows density of ADP in the P1b-state, ADP or mixed ADP/ATP occupancy in the P1b-state and ATP in the P1c-state.

Transition from the P1c- to the P2a-state would require seam subunit 6 to adopt the T-conformation, whilst the subunits 4 and 5 switch to the D-conformation and the S-conformation, respectively. This rearrangement would allow the γ-phosphate of subunit 6 to interact with the R652-finger of the subunit 1, stabilizing the A-domain of the subunit 6 in the T-conformation. The pore-loop1 of subunit 6 is located at the bottom of the A-tunnel. Here, its YV-pincer grabs the extended substrate protein chain, and its A-domain takes on the T-conformation and an orientation that is very similar to the A-domain of subunit 1 in the P1-states. These structural rearrangements result in an upward movement of the pore-loop1 motifs of subunits 1 to 4 relative to the N-domain assembly, pulling up the extended substrate protein chain grabbed by their YV-pincers. Since neither ATP hydrolysis nor ADP/ATP exchange appear to occur in the P1c-to-P2a transition, we propose that partial unfolding of the substrate protein is the rate limiting step.

A similar set of conversions reverts the P2-state to the P1-state, but now subunits 3 and 4 are the seam subunits. This transition has a critical implication for the orientation of the N-domain assembly. Because the orientation of the N-domain assembly relative to the A-domains is linked to the position of seam subunits in the P1a-state, the threefold axis of the N-domain assembly must tilt from subunit 4 towards subunit 2. Thus, the mechanism we propose is characterized by a sixfold cyclical transition of A-domain conformations and a threefold cyclical interconversion of P1- and P2-states.

The proposed mechanism may explain the structural variability of the N-domain assembly, that is witnessed by its relatively poor resolution (fig. S2 and S4). Our data indicate that the N-domain assembly binds the folded part of the substrate protein, and that its extended terminal strand is threaded thought the N-cavity and A-tunnel, where it is held (fig. 3C, fig. 5). Thermally driven large-amplitude motions of the N-domains would subject the folded part of the substrate protein to a wide spectrum of pulling forces. This would increase the probability that residues preceding the extended substrate protein strand unfold. Subsequent relaxation of the N-domain assembly would reduce the tension on the unfolded strand. This would either allow refolding or the A-domains to reel in the slack by means of the proposed catalytic cycle.

We propose that LonP1 functions as a Brownian, or Feynman–Smoluchowski ratchet (38, 39). Such a device uses random, thermal fluctuations for generating unidirectional motion or force, by impeding the reverse fluctuation, very much like a pawl blocks a ratchet from rotating backwards. Feynman showed that blocking this reversal requires energy. In LonP1, reversal would be blocked by the conformational changes that follow ATP hydrolysis. ATP hydrolysis upon substrate protein translocation must be fast compared to other catalytic steps, since the P1a- and P2a-states that we propose immediately follow translocation, could only be observed in the presence of the slowly hydrolyzing ATPγS. We propose that ATP hydrolysis is triggered by substrate protein translocation and that other Lon proteases share this mechanism. Our data indicate that ADP/ATP exchange by the first seam subunit occurs prior to protein substrate translocation in the transit from the P1a- to the P1c-state and from the P2a- to the P2c-state. However, our data do not establish whether this ADP/ATP exchange speeds up unfolding through a power stroke. We must consider this possibility since a ratchet mechanism does not exclude a power stroke. However, a power stroke may not be required if the observed flexibility of the N-domains would be amplifying the effect of thermal vibrations on unfolding of the substrate protein.

LonP1’s many active states would allow controlling its activity by effectors like specific DNA or RNA sequences, that could act by stabilizing or inducing specific states and/or rigidify the N-domains. Such processes may be essential for explaining why and how LonP1 can fulfill its role as a central hub in regulating mitochondrial health.

## Materials and Methods

### Cloning, protein isolation and purification

The human LONP1 gene (wild-type) and LONP1 mutants (K529R, E591Q, E801A, T803V, E812A, H841F, H843F and S855A) after the signal sequence (amino acids 125 to 959) were codon optimized and cloned into a pET11a expression vector with an N-terminal hexa-histidine tag, leaving two extra GS residues between the tag and the M125 residue (GenScript). Wild-type and the associated LonP1 mutants were expressed in BL21 (DE3) *E. Coli* strain that was cultured in 2XYT medium at 37°C and was induced overnight with 0.4 – 1.0 mM IPTG at 18°C. All the subsequent steps of purification were done in batch mode at 4°C. The filtered cellular lysate (in buffer A: 50 mM HEPES, 300 mM NaCl, 10 mM MgCl_2_, 20 mM imidazole, 10% (v/v) glycerol, 2 mM β-mercaptoethanol, Roche-protease inhibitor cocktail, pH 7.5) after sonication and centrifugation was incubated with Ni-NTA slurry for 60 minutes or overnight for binding. Sequential washes with high salt buffer (buffer A with 0.5 M NaCl, no imidazole) followed by imidazole buffer (50 mM HEPES, 150 mM NaCl, 10 mM MgCl_2_, 40 mM imidazole, 10% (v/v) glycerol, 2 mM β-mercaptoethanol) were performed to reduce contaminants. The protein was eluted with a step gradient of imidazole from 0.2 M to 0.4 M imidazole in buffer A containing 150 mM NaCl. Based on the purity from SDS-PAGE, the eluted fractions were subsequently passed through a QFF-Anion exchange column, or directly on the size exclusion Superose 6 Increase 10/300 GL column (figure S12). The pure fractions were pooled, snap-frozen and stored at -80°C for later functional and biochemical studies. To limit aggregation during cryo-EM grid preparation, LonP1-grids were prepared on the same day without freeze-thaw cycle.

The specific substrate protein of Lon, TFAM (Addgene plasmid #34705, Human_TFAM_NoMTS_pET28 was a gift from David Chan) was expressed and purified following a published protocol (8). The protein was stored in 10% (v/v) glycerol at -80°C.

### ATPase assay

The ATPase activity of wild-type LonP1 and its mutants was measured with ATPase/GTPase activity assay kit (Sigma-Aldrich, USA, product: MAK113) as per the manufacturer’s protocol. The reaction mixture (containing 100 nM protease/mutant enzyme; either 2 or 100 nM FITC-casein or 100 nM TFAM protein substrate and 1 mM ATP) was incubated in the ATPase buffer (20 mM HEPES, 100 mM NaCl, 10 mM MgCl_2_, 0,1 mM EDTA, 1 mM β-mercaptoethanol, 10 % glycerol, pH 7.8) for 30 min at room temperature, and then the reaction was stopped with the addition of malachite green. Finally, the absorbance at 620 nm was measured after a waiting time of 30 min with a Tecan Spark^®^ Multimode Plate-reader. All the measurements were performed in triplicate (three independent measurements) in 96-well plates for the accurate analysis.

### Protease assay

The proteolytic activity was analyzed with a constant amount of 2 nM of FITC-casein as protein substrate in the reaction mixture containing wild-type LonP1 or the corresponding mutants (100 nM) and ATP (1 mM). All the measurements were done in triplicates (from three independent assays) at RT in 96-well microplates for fluorescence-based assays (Invitrogen, REF M33089).

### Mass spectrometry

MS was employed as a sensitive tool to accurately determine the proteolytic activity of LonP1 and its mutants *in vitro.* Essentially, 1 µM of the wild-type and protease mutant protein were incubated with 1 µM of FITC-casein for 3 hours at room temperature in the reaction buffer (20 mM HEPES, 100 mM NaCl, 10 mM MgCl_2_, 0,1 mM EDTA, 1 mM β-mercaptoethanol, 10 % glycerol, pH 7.8, 5 mM ATP or 5 mM ADP). The end-products (peptides) of the reaction were separated through a 30 kDa cut-off filter (Amicon® Ultra-0.5 mL Ultracel®) to collect the flow-through and were mixed with 1 µL of 5 % TFA. Finally, the peptides were subjected to mass spectrometry and analyzed using Mascot (Matrix Science, London, UK; version 2.4.1) assuming a non-specific digestive enzyme. Protein probabilities were assigned by the Protein Prophet algorithm (40, 41) and were annotated with GO terms from NCBI (42). All the subsequent peptide coverage (in %) were stated in table S2. CLUSTALW (43) was used for the multiple sequence alignment of the human LonP1 and their related homologs.

### Grid preparation and data acquisition for cryo-electron microscopy

For datasets I and II, 3.5 µL of wild-type LonP1 (∼0.7 mg/ml in 20 mM HEPES, 100 mM NaCl, 10 mM MgCl2, 1 mM β-mercaptoethanol, pH 7.5) was applied onto glow discharged 300 mesh Lacey carbon grids. Prior to freezing, either 1 mM AMPPCP (dataset II) or 1 mM ADP-ATP mix (dataset I) was added, and the protein sample was incubated for 10 minutes on ice. The grids were plunge-frozen in liquid ethane after blotting away excess sample for 3 s under 100 % relative humidity conditions, using a Vitrobot IV (Thermo Fisher Scientific) that was maintained at 10°C. For dataset III, LonP1was mixed with TFAM at 1:2 molar ratio (final concentration ∼ 0.5 mg/ml) in the presence of 1 mM ATPγS for 5 min prior to Vitrobot freezing on 300 mesh R 2/2 Quantifoil copper grids (Electron Microscopy Sciences). All three datasets were collected on a FEI Titan Krios TEM with a GATAN post column energy filter (20 eV zero-loss filtering) operating at 300 kV.

For datasets I and II, super-resolution movies of 40 frames were acquired in microprobe mode using a Gatan K2 summit detector, whereas for dataset III, it was operated in counting mode (table 1). All the movies were acquired using the SerialEM software package and were then preprocessed for pruning in FOCUS (44) during data acquisition. Briefly, the super-resolution movies were clipped and binned 2X with the aid of IMOD and Frealign packages respectively. MotionCor2 (45) was used to align and correct the frames with an approximate dose weighting. The contrast transfer function (CTF) was estimated with CTFFIND4 (46) on the non-dose weighted aligned movie frames. Micrographs with lower CTF estimation (less than 7 Å) values, large beam-induced drifts and crystalline ice were excluded from further single particle analysis (47).

### Image processing and map generation

For dataset I, the CTF defocus parameters of aligned averages from the movies were once again estimated with Gctf (Gctf_v1.06 and above). Roughly, 20,000 reference-free particles (from 400 preselected micrographs) were picked with Gautomatch (K. Zhang, MRC LMB; https://www2.mrc-lmb.cam.ac.uk/research/locally-developed-software/zhang-software) to perform three rounds of 2D classification in Relion (48–51). The best and well centered 2D averages were then low pass filtered to 20 Å and used as templates to pick the particles from the entire aligned averages that resulted in roughly 400,000 particles. After 4 rounds of 2D classification, ∼280,000 particles were subjected to 3 rounds of 3D classification with either K=4 or 6 or 8 respectively. Two distinct classes (1, 2) and (3, 4) from the K=6 3D classification were selected based on the closing of the gap between the subunits. These were subjected to refinement and postprocessing in Relion. Particles belonging to these classes 1 and 2 were processed again in CryoSPARC (52) with additional selection and pruning to exclude the bad particles. A total of 89,000 particles belonging to the open class were subjected to a final local-refinement in CryoSparc to yield an R-sate at 3.7 Å resolution (FSC at 0.143 gold standard). However, the local refinement of closed classes (120,000 particles) did not show any improvement in the resolution or the map quality. The combined closed class1-2 particles were subjected to an additional focused 3D-classification (with Tau=10 and K=4) in Relion with a loose mask around the A- and protease domains, excluding the flexible N-domain. This resulted in three good classes with subtle differences either in the A- or A- and N-domains, and were named as P1b-, P1c- and P2-states. To improve the resolution and map quality, these states were then subjected to non-uniform and local refinements in CryoSPARC to yield maps of P1b-, P1c- and P2-states at 3.9, 4.3 and 5.8 Å respectively.

For Dataset II: 2D classes from dataset I were used as templates to pick particles that were subjected two rounds of 2D-classification in Relion, yielding 320,000 particles. After two rounds of 3D-classification with K=4, particles belonging to class2 (∼ 126,000 particles) were selected for refinement and post-processing in Relion, resulting in a 4.23 Å R-state map (FSC at 0.143). Particles from this R-state were once again pruned and selected in CryoSPARC after heterogenous refinement, that resulted in a clean R-state (with 76,000 particles). Local refinement of this R-state improved the map to 3.75 Å (FSC at 0.143).

For Dataset III: Following the same strategy as mentioned above for dataset II, a single processing state, P1a (78,000 particles) was obtained after a final refinement and postprocessing in Relion. This P1a-state was further subjected to focused classification (K=3) with a loose mask around the rigid A- and protease domain to yield a better resolved P1a-state (3.8 Å) and a low resolution mixed-state map. Heterogeneous refinement in CryoSPARC with K=3 classes yielded the low-resolution docking states D1 and D2, and the P2a-state.

The difference between the masked and unmasked FSC curves was mainly caused by the large box size we used (FSC curves in fig. S2, S3 & S4). Masking out the volume outside the LonP1 density therefore reduces the noise significantly, explaining why masking improved the FSC curves. Since LonP1 has an elongated structure, a spherical mask still encloses a substantial part of the box that does not correspond to LonP1 density, explaining the relatively limited improvement of the FSC curves upon imposing a spherical mask. All masks included the N-domains, which had a lower local resolution than the C-terminal domains. This explains the bimodal drop observed in several FSC curves: the first drop reflects the lower resolution of the N-domain density, the second drop corresponds to the C-domains.

### Model building, refinement, and analysis

Models were built and refined using Coot (53), ChimeraX (54), and Phenix-1.18 (55). Real space refinement included hydrogens to reduce potential clashes. Movies were calculated using ChimeraX, by interpolating between states.

## Acknowledgements

We thank Henning Stahlberg, Mohamed Chami, Lubomir Kovacik and Kenneth Goldie for supporting cryo-EM data collection at the Biozentrum BIO-EM lab; Rob McCleod, Ricardo Righetto and Ricardo Adaixo for advice on data processing; all C-CINA members for advice; SciCORE (http://scicore.unibas.ch/) scientific IT service of University of Basel for supporting Cryo-EM image analysis; Timothy Sharpe and Alexander Schmidt for the biophysics and MS facility; David C. Chan for providing a TFAM-encoding plasmid.

## Author contributions

IM expressed and purified LonP1 and TFAM, determined its 3D structure by cryo-EM, built the initial models, refined, and interpreted the structures and wrote the first draft of the manuscript. KS purified protein and its mutants performed protease and ATPase assays and wrote the manuscript. NS purified protein and its mutants and performed protease and ATPase assays. AT analyzed the final models and wrote the manuscript. TM advised on initial model building, refinement and on writing the manuscript. JPA supervised the project, built the initial models, refined, and interpreted the structures and wrote the manuscript.

## Data and materials availability

Models and maps have been deposited at the protein data bank: 7NGL (R-state in the presence of ADP and ATP), 7OXO (R-state in the presence of AMPPCP), 7NGP (D1-state), 7NGQ (D2-state), 7NFY (P1a-state), 7NG4 (P1b-state), 7NG5 (P1c-state), 7NGC (P2a-state), 7NGF (P2c-state). Subunits 1-7 are labelled as A-F in all coordinate files. All other data are available in the main text or the supplementary materials.

## Movies

**Movie M1. LonP1 binding substrate protein and going through its catalytic cycle.** Odd-numbered subunits are in in red, even subunits in yellow, the substrate protein is in green. The atoms of the substrate protein and the YV-pincers are indicated as spheres.

**Movie M2. YV-pincers grabbing, pulling, and releasing the extended substrate protein chain in the A-tunnel.**

## Supplementary Materials for

**Table S1.**
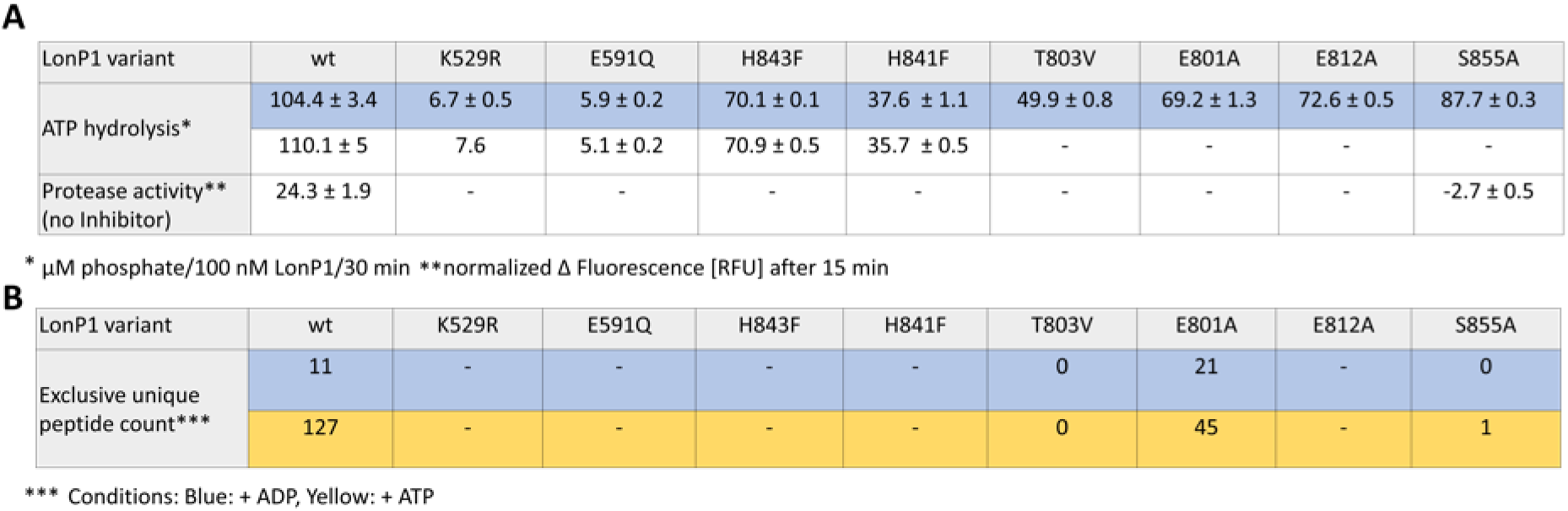
Enzymatic activity of LonP1 and LonP1 mutants. (A) ATP hydrolysis and protease activity for LonP1 variants. No substrate (blue background) or 100 nM TFAM (white background) was added to the ATP hydrolysis experiment. The protease activity was measured for LonP1 wt and S855A without added Inhibitor during the purification. (B) The peptide count (LonP1 peptides) of the mass spectrometry experiment is given (blue background: ADP was added, yellow background: ATP was added).

**Table S2.** Protease specificity of LonP1 and LonP1 mutants. Percentage (%) of amino acids identified from LonP1 and β-casein from the mass spectrometry experiment. Blue, yellow and red backgrounds indicate percentages from LonP1 peptides, white backgrounds from β-casein peptides.

| LonP1 variant | wt | T803V | E801A | S855A |
| --- | --- | --- | --- | --- |
| [%] of amino acids identified*** | 4.3 | 0 | 9.7 | 0 |
|  | 56 | 0 | 25 | 1.3 |
|  | 53 | 3.2 | 23 | 0 |
|  | 62 | 0 | 68 | 0 |

\*\*\* Conditions: Blue: + ADP, Yellow: + ATP, Red (LonP1 peptides): + ATP, FITC-casein, White (β-casein peptides): + ATP, FITC-casein

**Figure S1.**
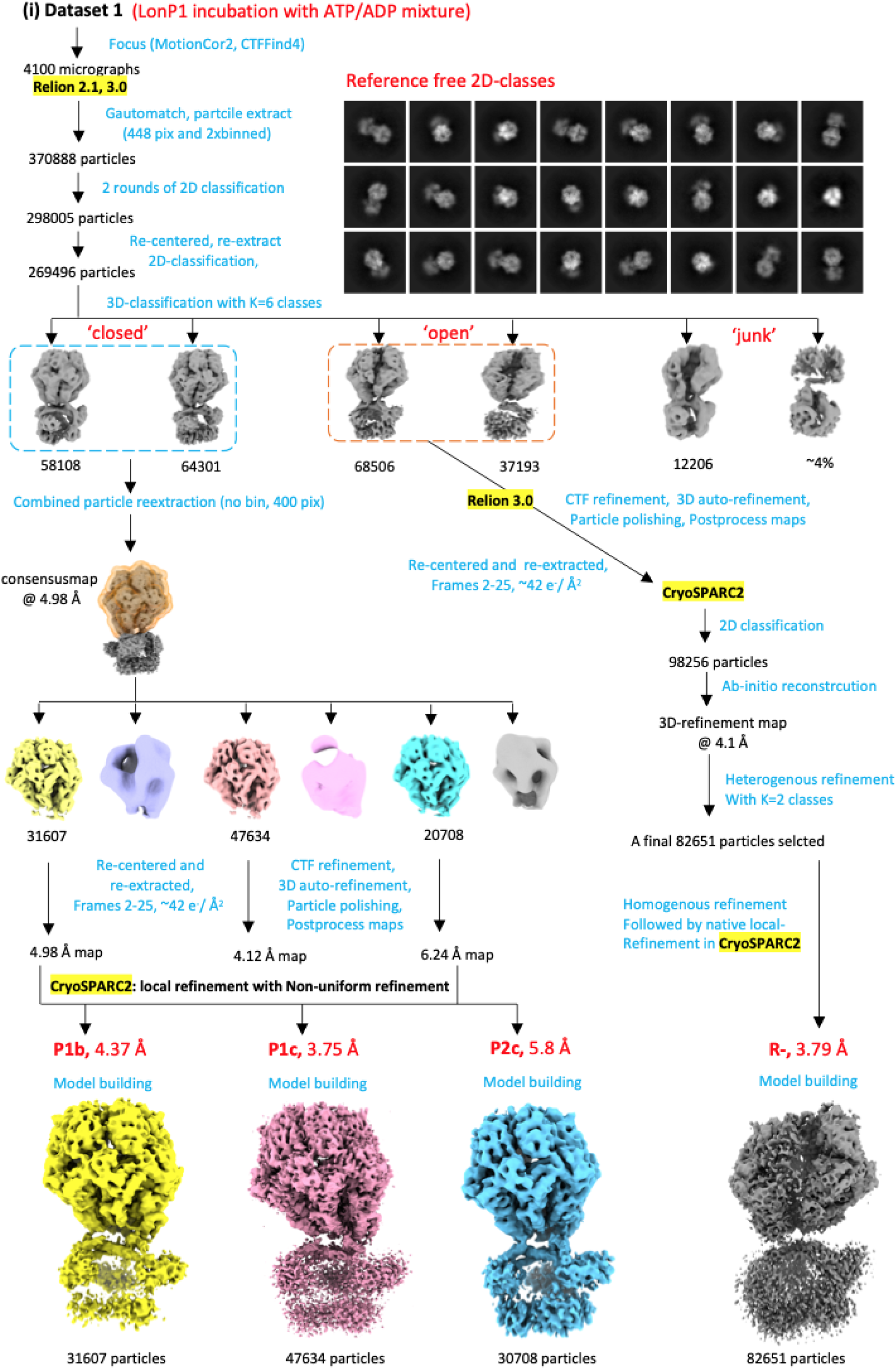
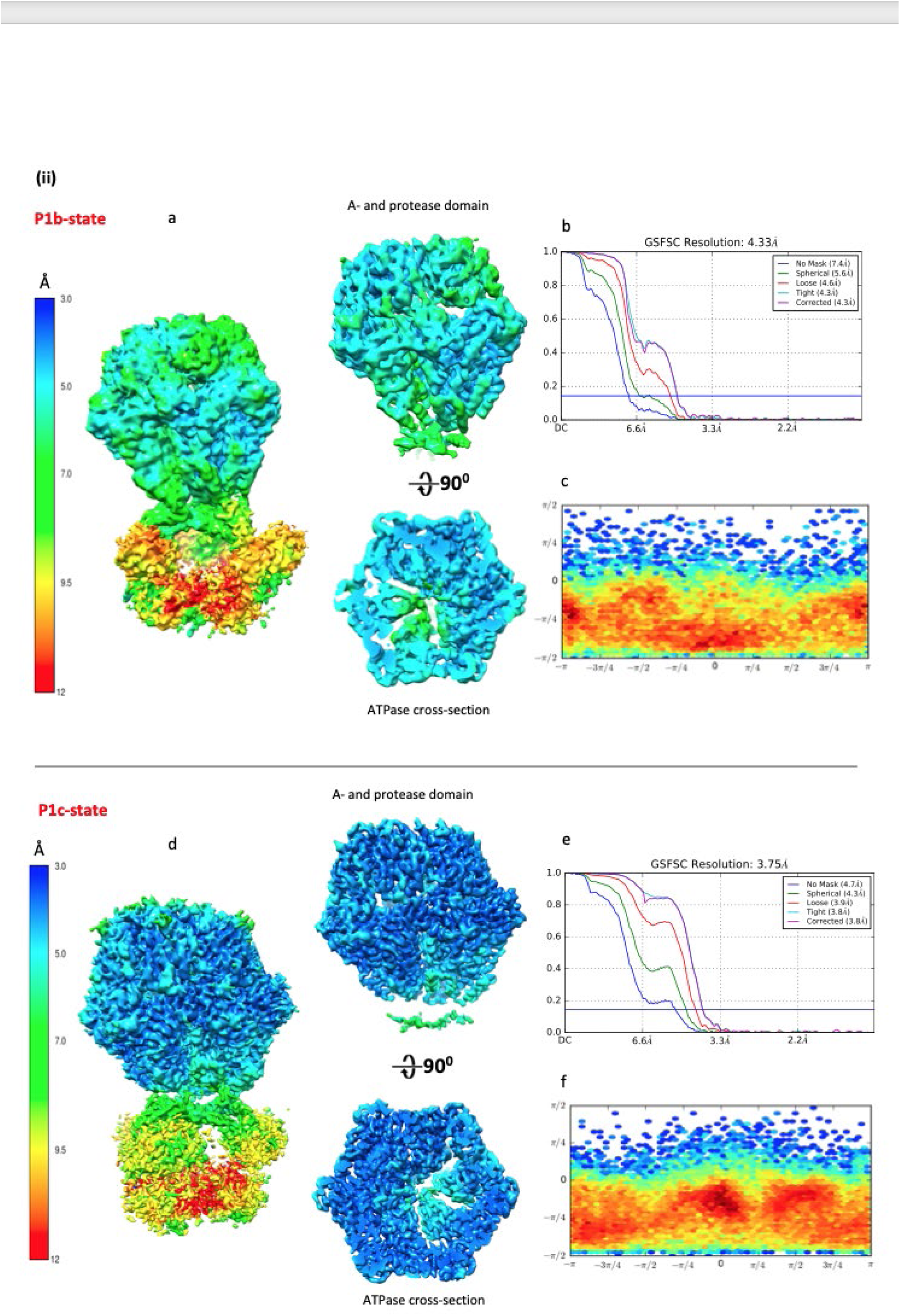
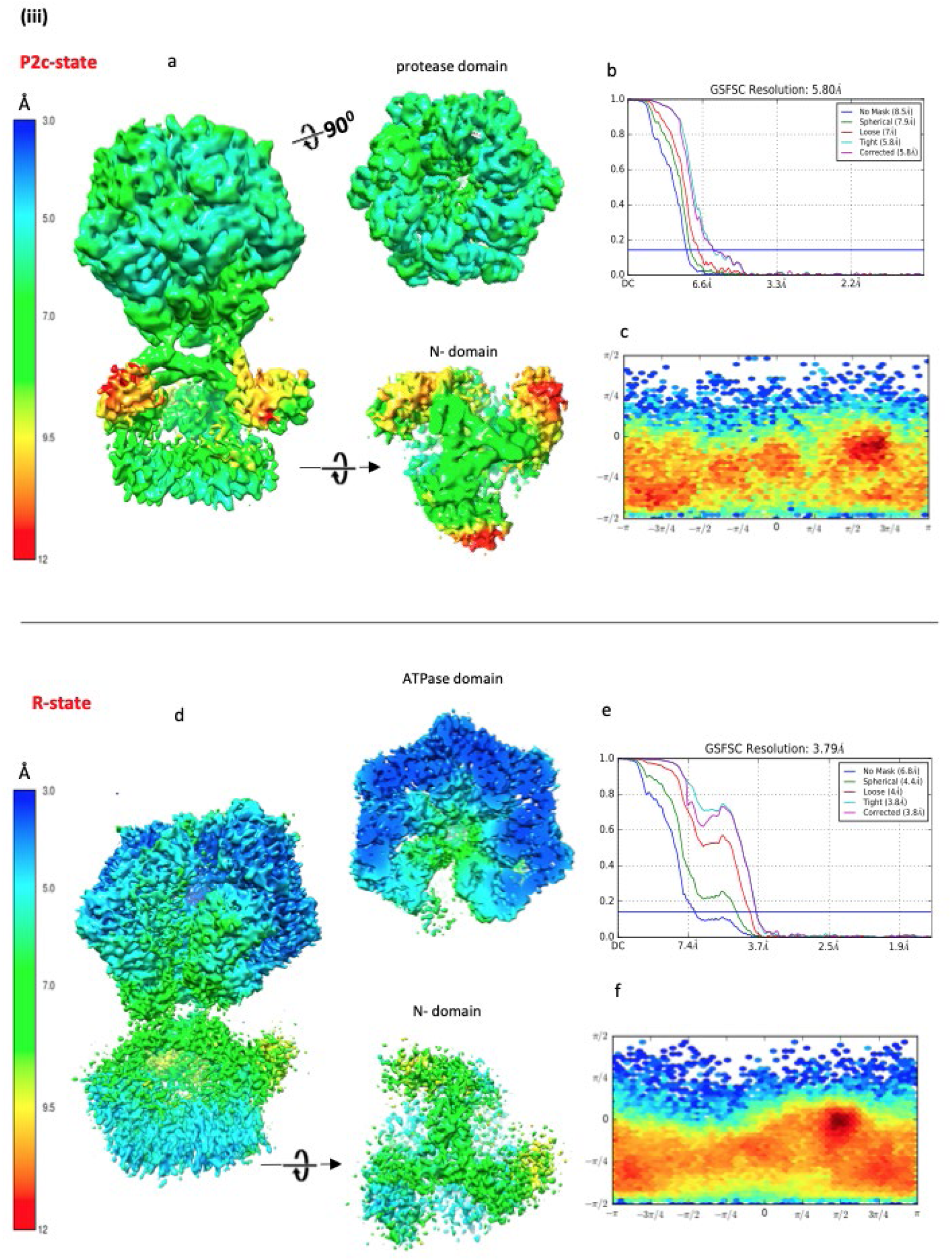
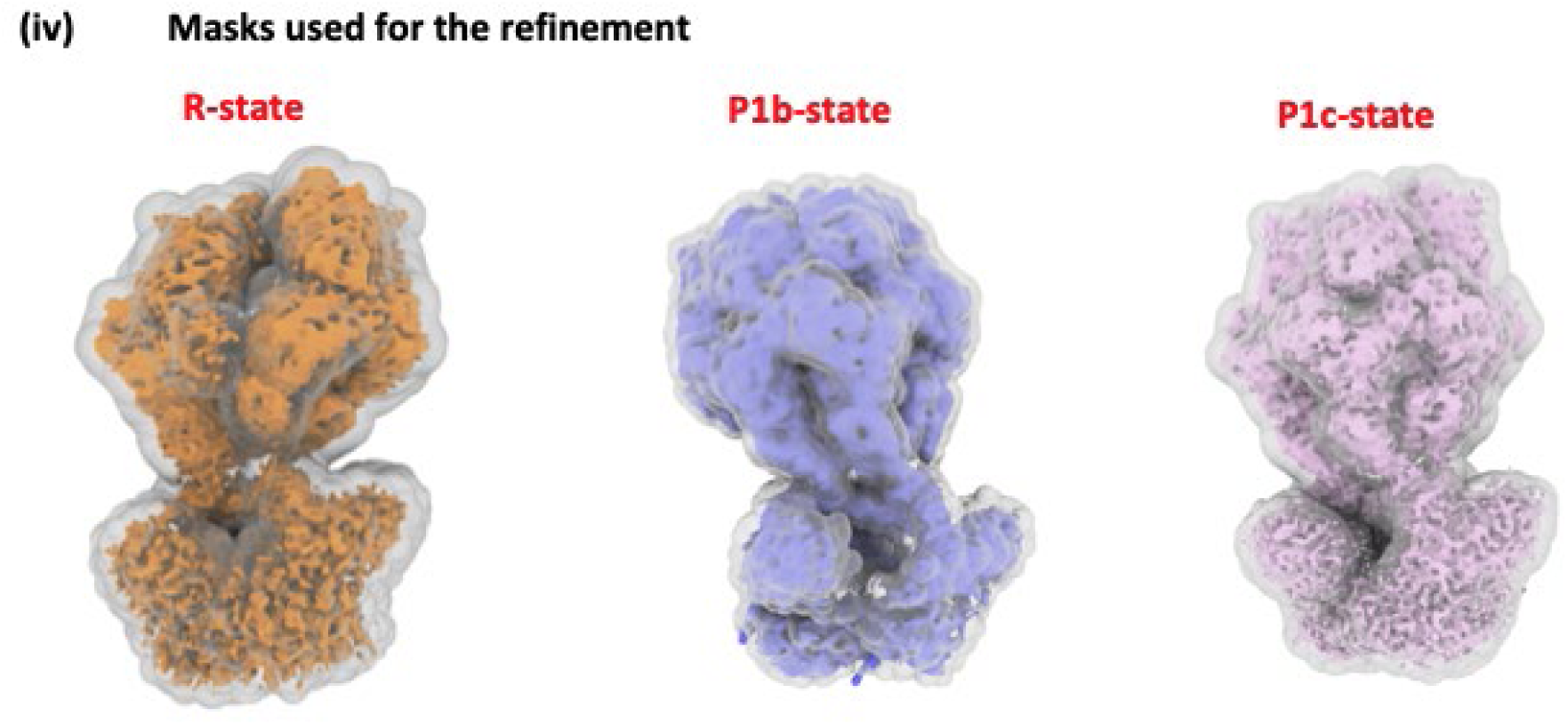
Data collection and 3D density calculation of LonP1 in ADP/ATP. (i) Summary of the results obtained from dataset I, resulting in the R-, P1b-, P1c and P2c-states. A total of 4100 movies were resulted in ∼370,000 full length LonP1 particles. After few rounds of 2D classification and pruning in Relion 2.1, a total of ∼270,000 particles were subjected to 3D classification. A sub-panel of 2D-class averages clearly show a threefold N-domain, as indicated with a red arrow. Two distinct classes named as ‘closed’ and ‘open’ were refined in Relion 3.0 and CryoSPARC2. The combined ‘closed’ particles after re-extraction and a final consensus refined map, yielded further three different maps (in yellow, P1b; pale pink, P1c and cyan, P2c colors) after an additional 3D classification without alignment, with a soft mask around A- and protease domain. (The A- and protease soft mask was in pale brown). The map resolutions and the quality were further improved after iterative CTF refinements and particle polishing and from the average of 2-25 movie frames (∼42 e/A^2^). The final maps from this dataset-1 were locally refined in CryoSPARC2 to build models in P1b- (31607 particles), P1c- (47634 particles), P2c- (30708 particles) and R-states (82651 particles) respectively. (ii and iii) The local resolution maps (a) of P1b-, P1c-, P2c- and R-states with their respective angular distribution (c) and FSC curves (b). The N-domain in all the states has a resolution range from 8-12 Å resulting from the continuous flexibility. (iv) The final soft-masks applied during the local refinement of R-, P1b- and P1c-states that included the N-domain.

**Figure S2.**
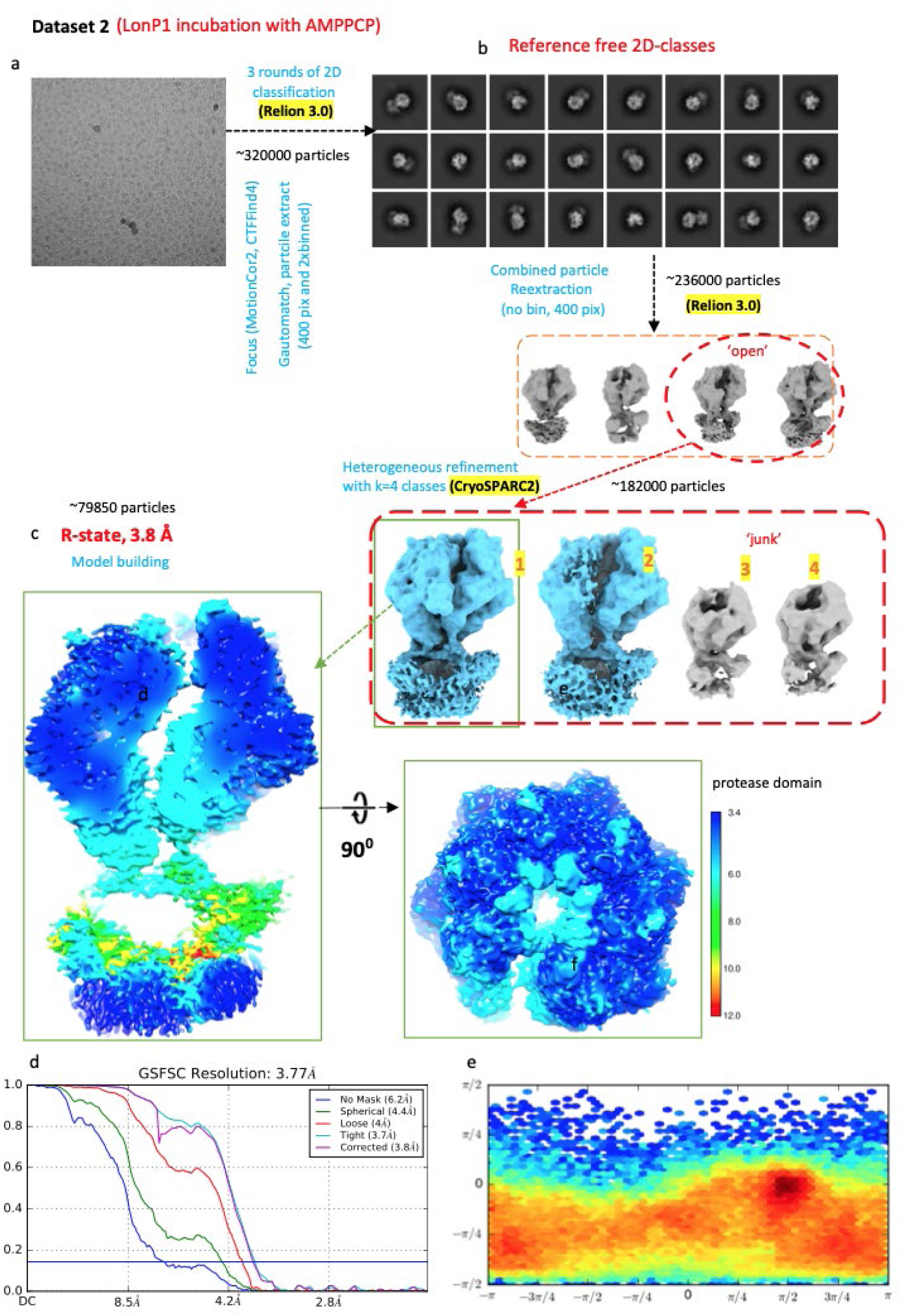
Data collection and 3D density calculation of LonP1 in AMPPCP. Summary of the results obtained from dataset II, resulting exclusively in the R-state after incubating with the AMPPCP. A total of 79,850 particles underwent in the final refinement of the R-state. The local resolution map (c) shows regions with 3.6 Å in the A- and protease domains and 10-12 Å at the N-domain. (d) and (e) are the FSC and angular distribution plots of this R-state.

**Figure S3.**
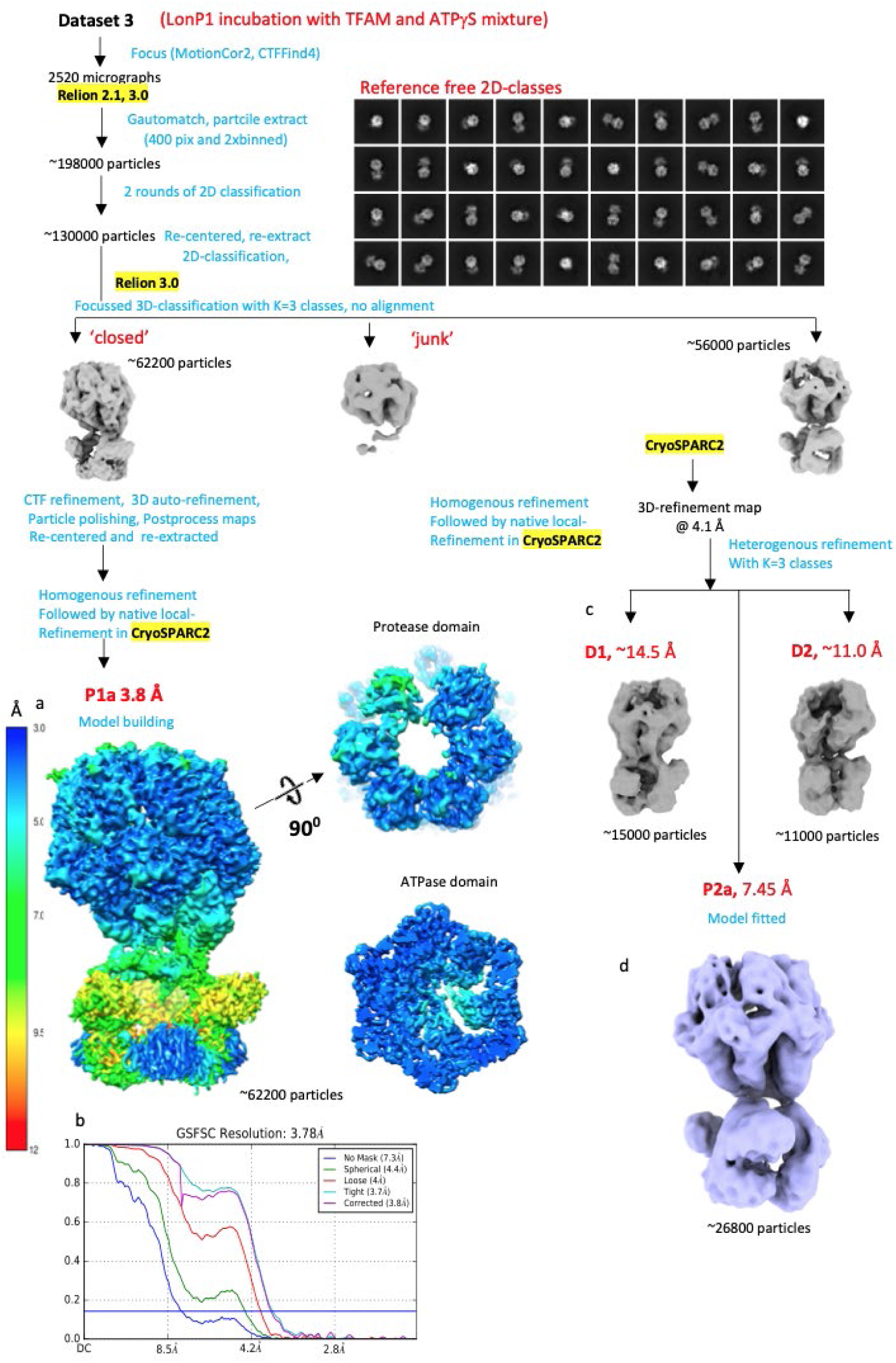
Data collection and 3D density calculation of LonP1 in ATPγS. Summary of the results obtained from dataset III, resulting exclusively in the P1a-state and a P2a-state (at lower resolution than in dataset II). In addition to the P-states, this incubation condition also favored to resolve the transient states (D1 and D2) of LonP1 in action during substrate recruitment (c).

**Figure S4.**
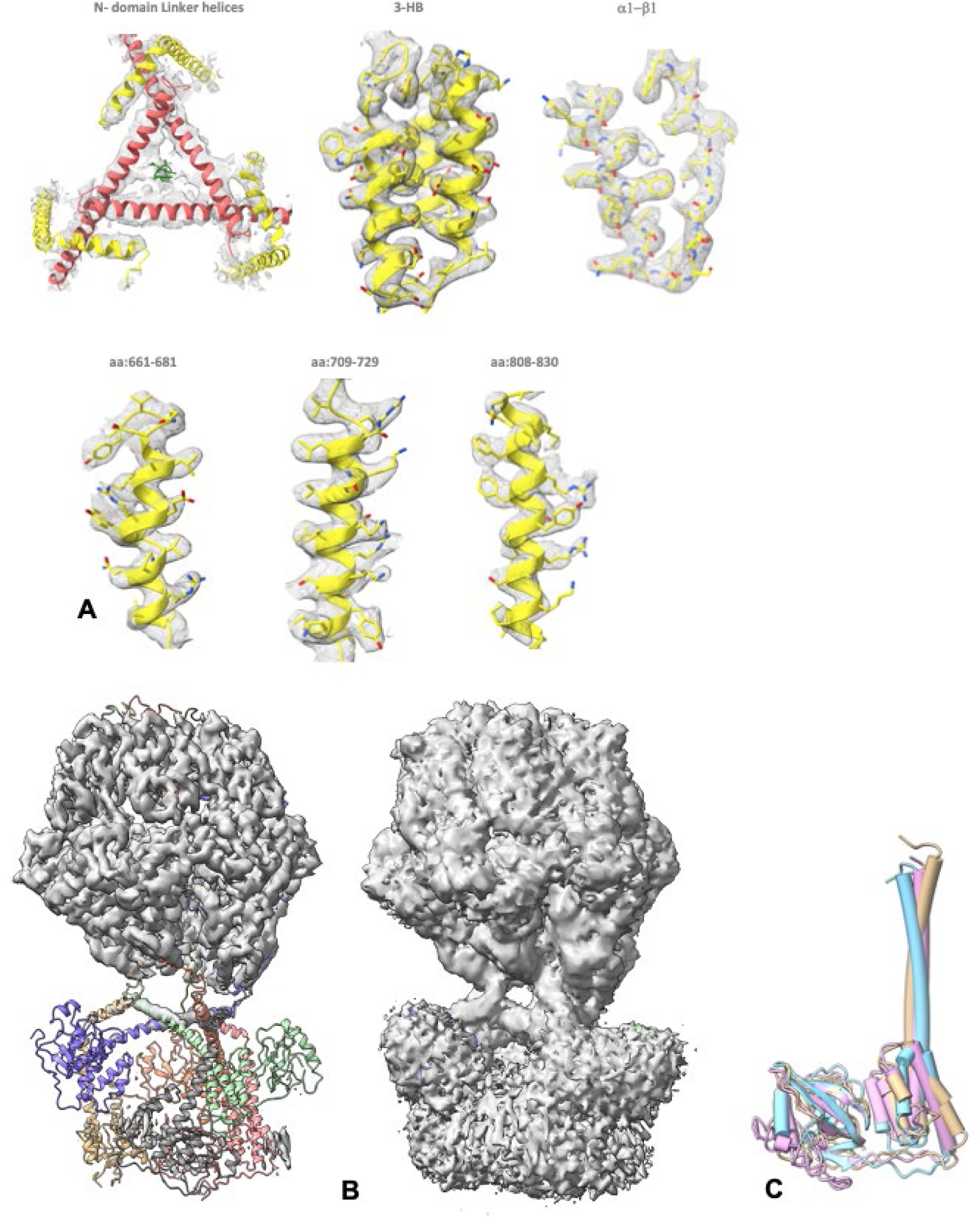
Fitting of the subdomain models and secondary structural elements into the EM density map. (A) The C-terminal α-helices of the N-domains could be recognized in the density, but not their side chains (top left). In better ordered parts of the maps, side chains could be localized (other panels). (B) Density of the N-domains: the P1a-state is shown at two different contour levels. A high contour level (left) reveals the well-resolved features, specifically the A- and protease domains. The atomic model is also displayed, showing that the N-subunits do not fit into the well-ordered density. However, when the contour level is reduced (right), also the more poorly ordered regions are revealed, and the location and orientation of the N-domain assembly becomes obvious. (C) Superimpositions of the structured parts of N-domain models generated by SWISSMODEL (orange), ITASSER (pink) and AlphaFold (cyan) show all models share the same fold and molecular shape.

**Figure S5.**
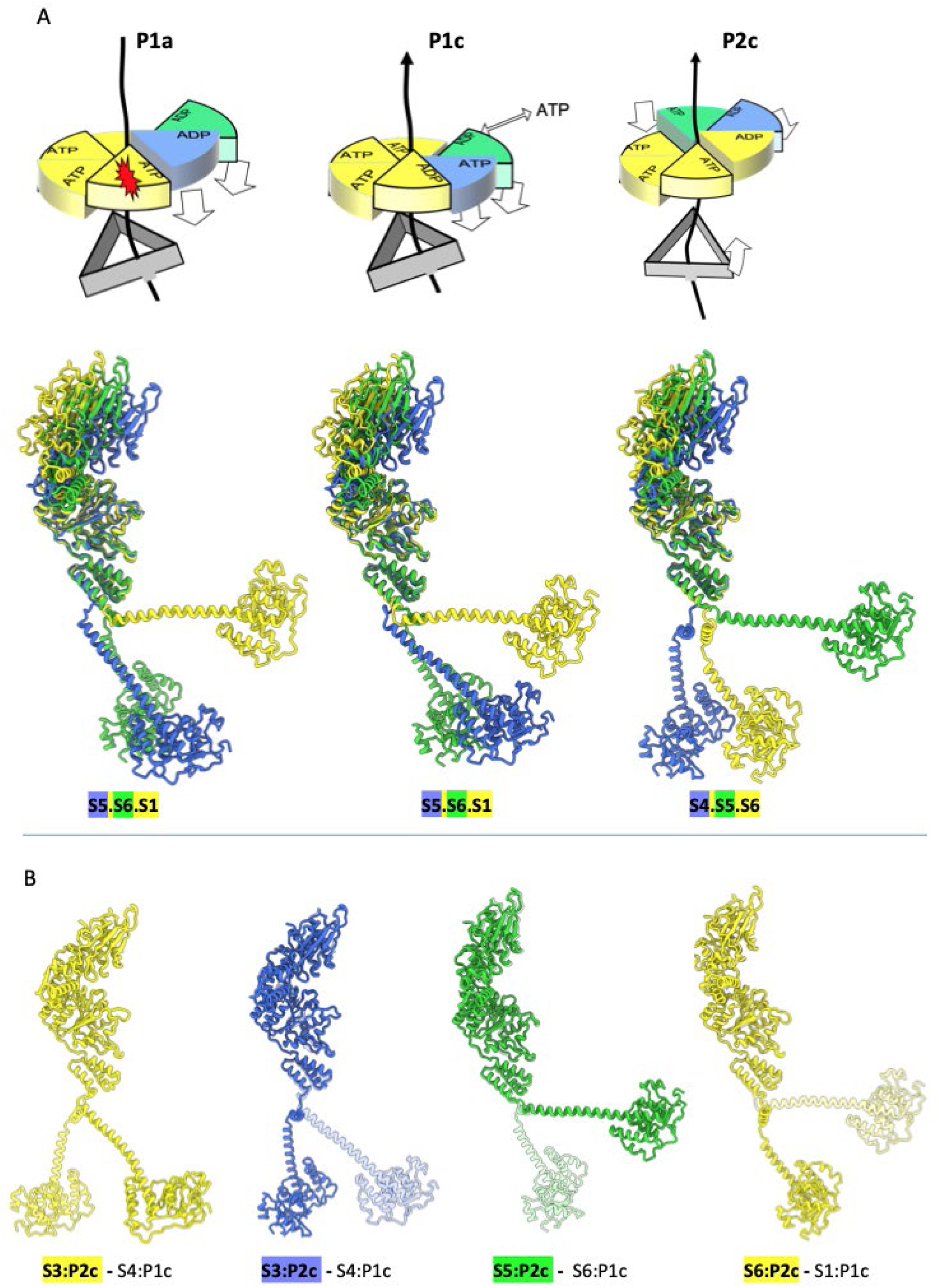
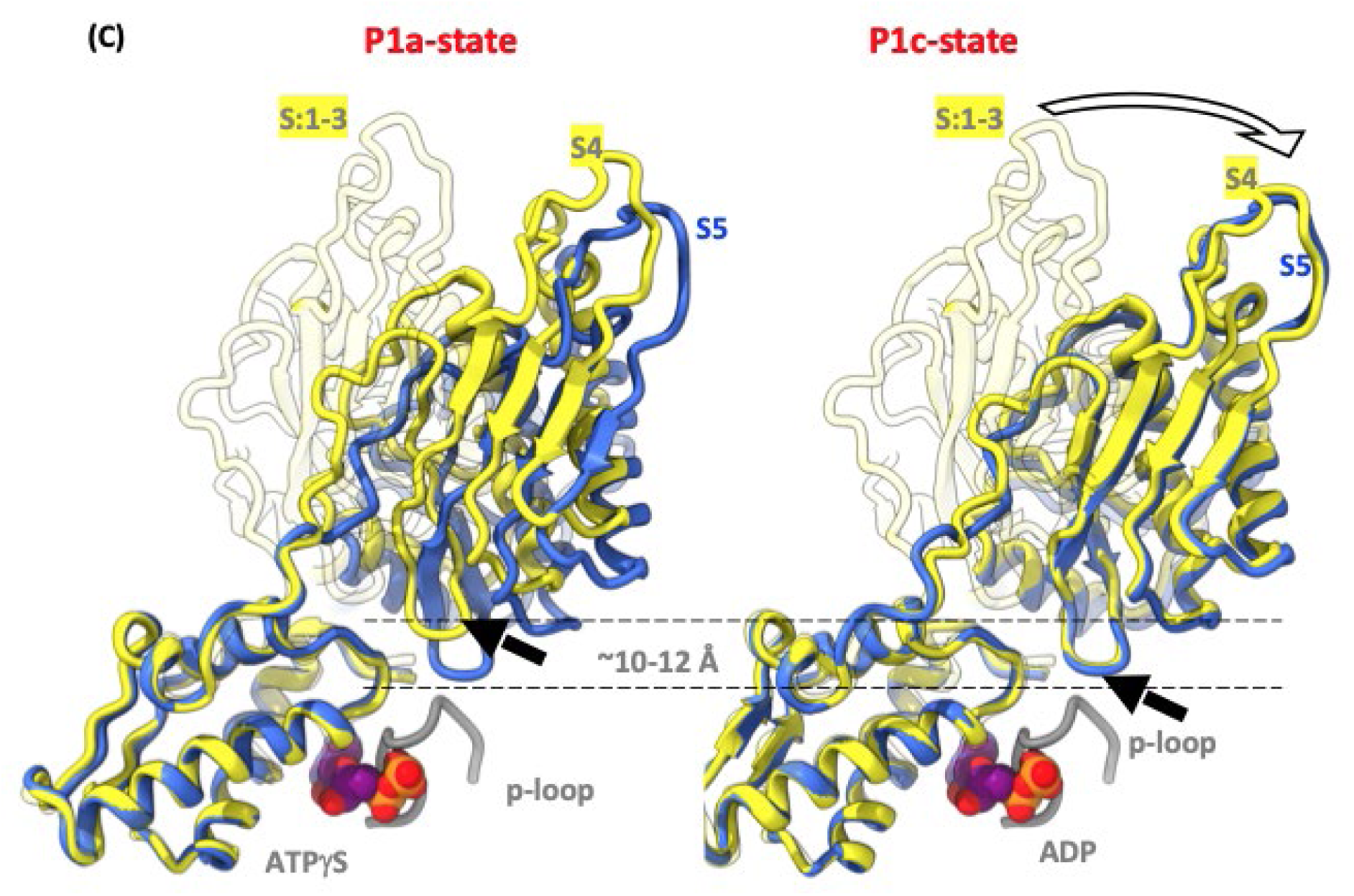
Rigid body movements of the LonP1 subdomains during the structural transition from P1- to P2-state. (A) Superimposition of the first seam subunit (in the D-conformation; blue), the second seam subunit (in green) and the first subunit (in the T-conformation; yellow). The first seam subunit in the P1a- and P1c sub-states is an odd (S1) subunit, whereas it is an even (S6) subunit in P2c. During structural transition from P1a- to P1c-substate, the protease domain of first seam subunit S5 (in blue) tilts by 20° along the pseudo threefold axis. (B) shows the perfect structural alignment of the protease and A-domains between the P1c- and P2c-substate, that were superimposed on their α-subdomains. The bold color representation depicts the subunits from P2c-substate and the light color show the corresponding subunits in P1c-substate. Except for orientation of the N-domain, the P2c-substate is very close match to the P1c-substate. (C) shows the zoomed view of the movement of protease domain during the ATP hydrolysis and exchange from P1a- to P1c-substate. The S5 seam subunit (in blue) of P1a-state (i.e., ADP bound) is in very close proximity to the P-loop of the nucleotide binding pocket. Whereas the S4 subunit (in yellow), which is still ATPγS bound in the P1a-state was 10-12 Å away. Upon ATP hydrolysis on the S4 subunit in P1c-substate, the β-hairpin loop of the protease domain moves close to the p-loop, as represented in the right panel (C). The orientations relative to the A-domains of the protease domains of seam subunits S5 of P1a-, S4 of P1c- and of the R-state subunits in the D-conformation differ less than 1 Å RMSD and are represented here in blue. The P1-state subunits S1 to S3 are depicted in light yellow color.

**Figure S6.**
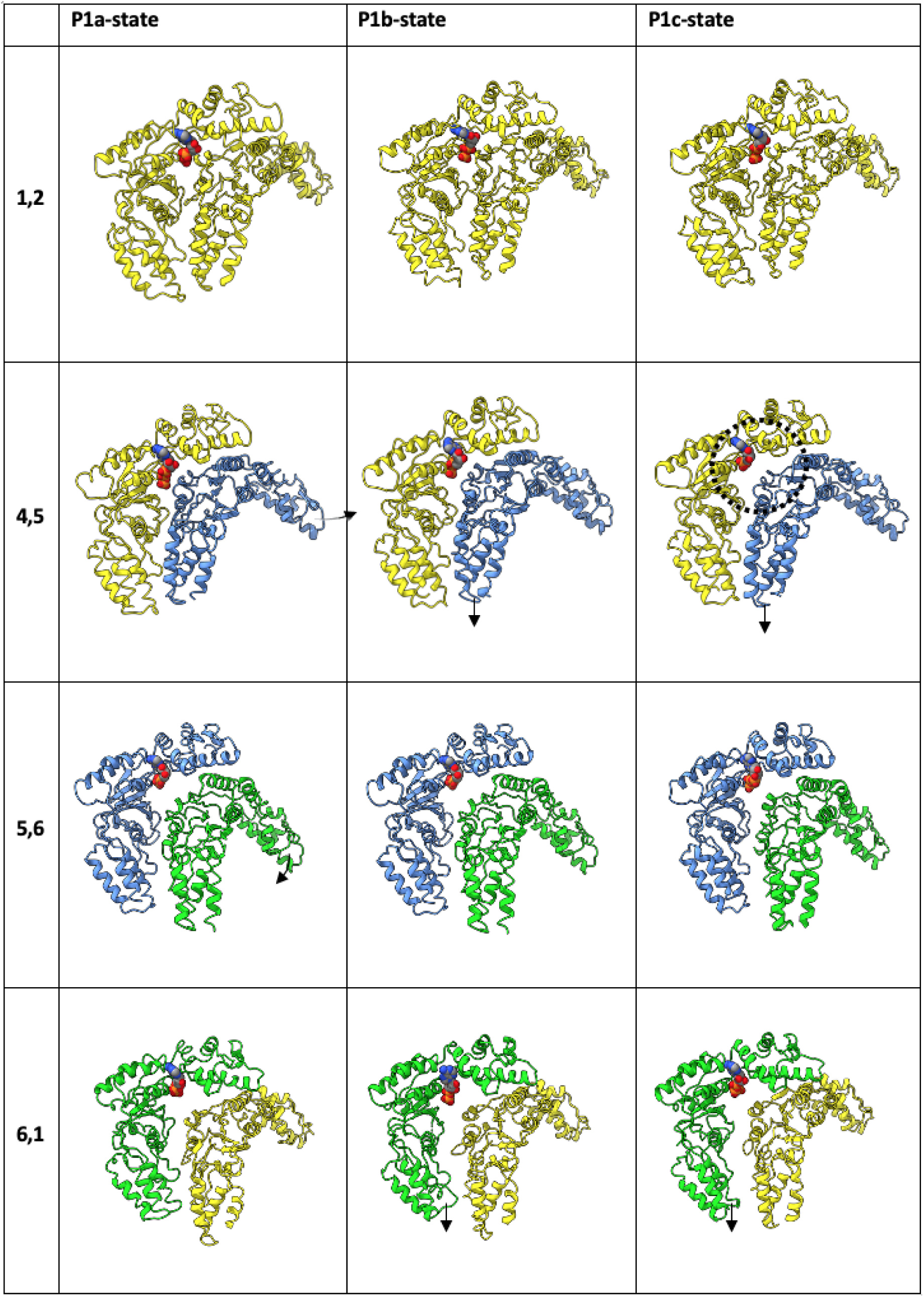
Interfaces between the A-domains of LonP1 as a function of processing state (columns) and subunit number (rows). The helical subunits (1–4) are in yellow and the two seam subunits 5 and 6 in blue and green, respectively. In each panel, the α*/*β-subdomain of the left-hand A-domain is an identical orientation towards the viewer, and its bound nucleotide is shown. Since the helical subunit pairs all have very similar interfaces, only the subunit 1/2 pair is shown for each of the P-states (cf. the first row). The outward tilt of the α-subdomain of the first seam subunit 5 compared to the interface of the helical subunits is indicated by a curved arrow in the P1a-state subunit 4/5 pair (column 1, row 2). The downward tilt in the second seam subunit 6 of the subunit 5/6 pair is also indicated by a curved arrow (column 1, row 3). The downwards motions of the seam subunits relative to the helical subunits are indicated by straight arrows in columns 2 and 3, rows 2 and 4. The release of the first seam subunit from the last helical subunit due to ATP hydrolysis and concomitant relaxation of the R652-finger in the P1c-state is indicated by a dashed circle (column 3, row 2). The interfaces of the helical subunits are commensurate with a right-handed helical rise (i.e., the subunit to the left is slightly lower than the righthand subunit). Interfaces that involve at least one seam subunit are unique and especially the 5-to-6- and 6-to-1 interfaces demonstrate a strong local left-handed helical rise to compensate for the righthand helical rise of the non-seam subunits (i.e., the subunit to the left is higher than the righthand subunit).

**Figure S7.**
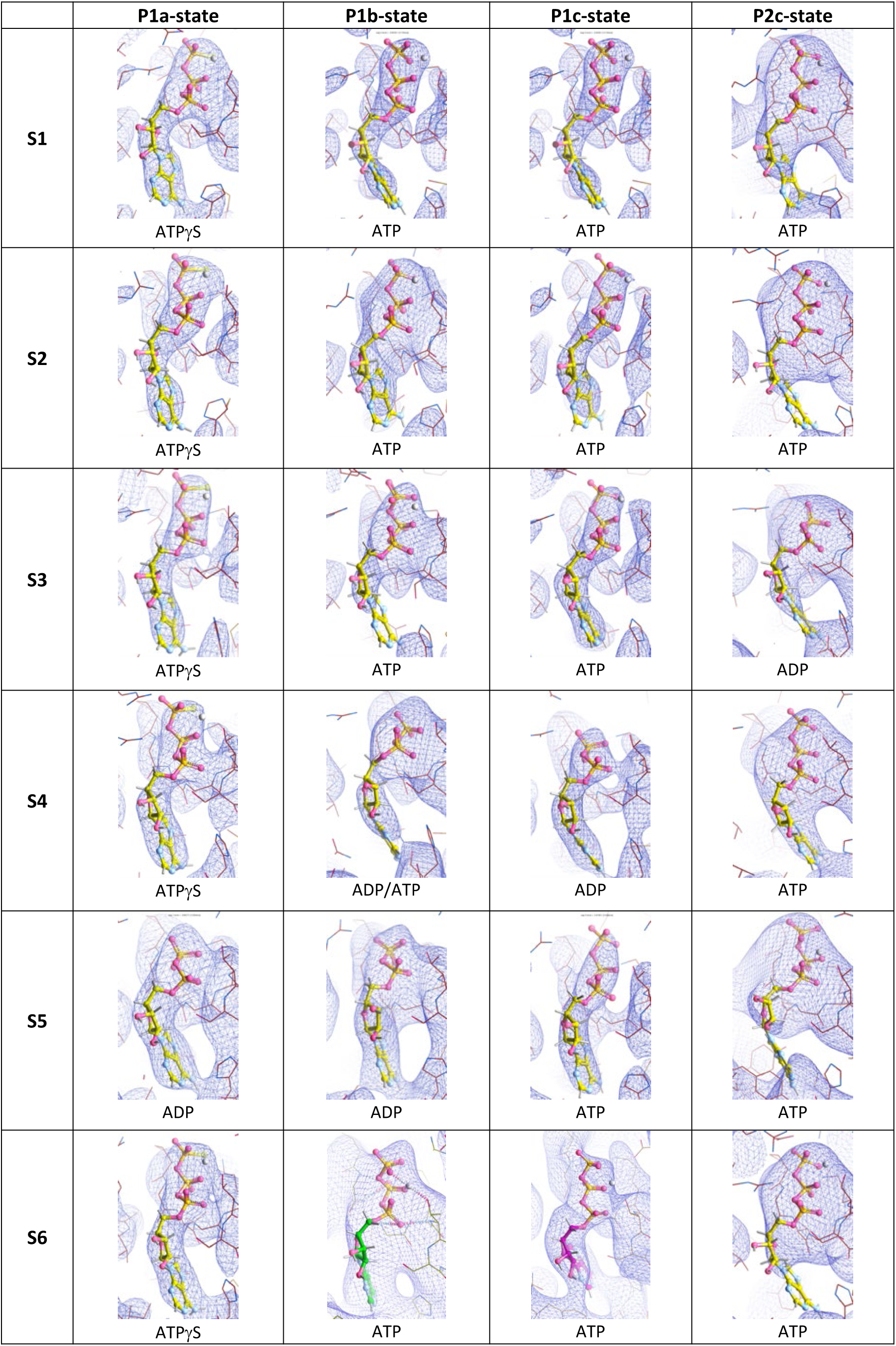
Nucleotides bound by subunits 1 to 6 in the P1a, P1b, P1c and P2c sub-states. The scattering potential is locally contoured at 2/3 of the observed scattering potential of either the Pa- or Pb- phosphorus atom of the bound nucleotide (whichever is lower). In most cases this gives a reasonable contour for the di- or triphosphate moiety of the nucleotide, but at lower resolution this criterion is not always unambiguous (e.g. a γ-phosphate would fall half-in, half-outside the contour level in the case of subunit 4 in the P1b-state). Therefore, also the proximity of local positive charges, especially of the trans-acting Arg652, were checked in optimally and consistently identifying the nucleotide. But we do not exclude the possibility of a mixed ADP/ATP occupancy for some of the binding sites. Note that in the P1-states, subunits 5 and 6 are the seam subunits, whereas in the P2-states, subunits 4 and 5 are the seam subunits. Also notice that in the case of the P1a-state nucleotide binding sites, the γ-phosphate is especially conspicuous because of the extra scattering potential of sulfur, since LonP1 was incubated with ATPγS for inducing this state. Also notice the relatively strong density often observed for ordered Mg^++^ in many of the ATP-bound states (but is strikingly absent in the binding site of subunit 5 in the P1c-state, despite clear density for ATP). The Mg^++^-ions are indicated as white spheres. Although this metal is relatively light, its high positive charge increases its electron scattering potential.

**Figure S8.**
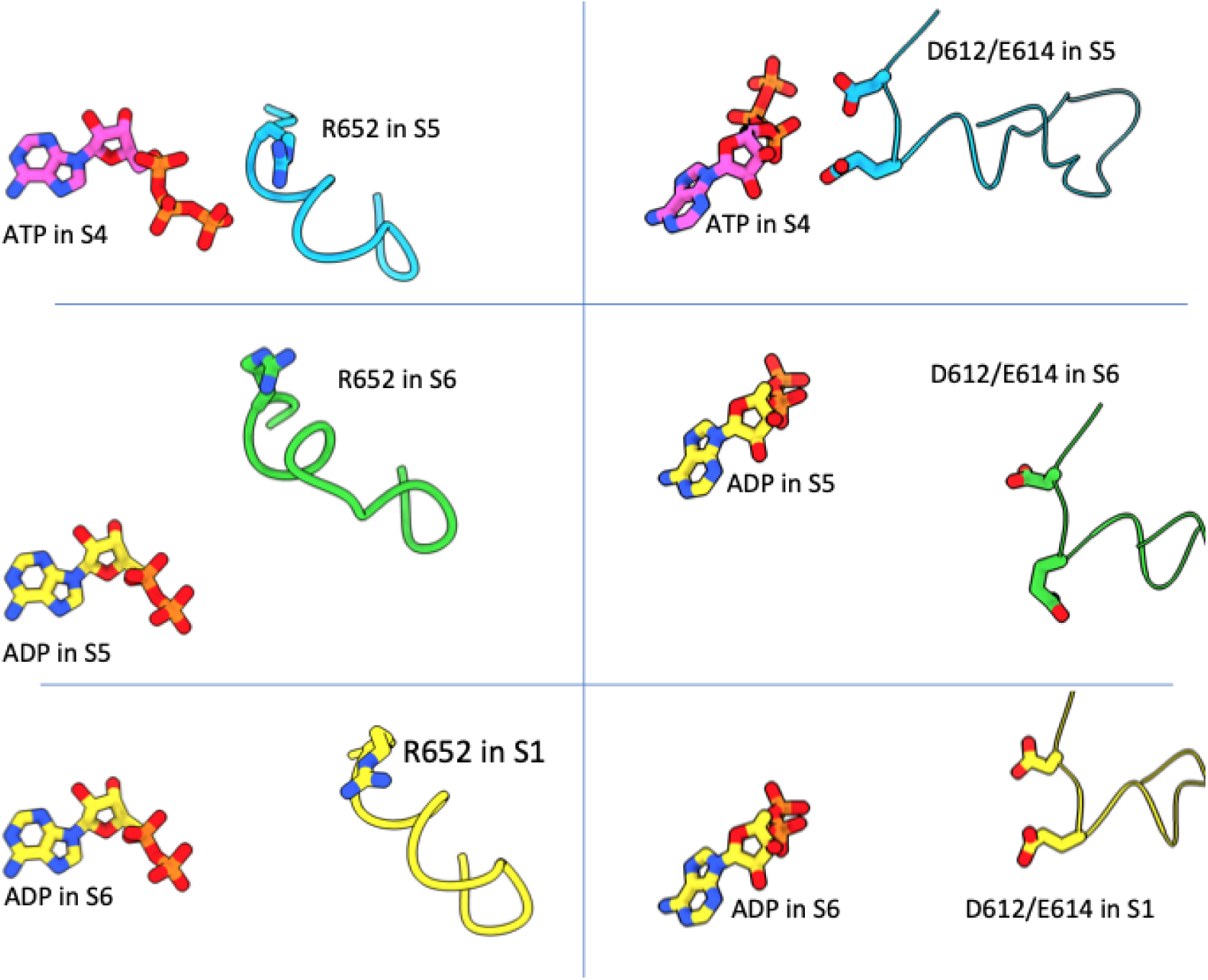
Trans-interactions with nucleotides in the P1a-state. The loop with the trans-acting R652-finger interact with the ATP γ-phosphate of the neighboring subunit (in P-conformation subunits, top-left panel), but when ADP is bound (as in seam subunits or in the R-state, lower left panels), this interaction does not occur. Only for D-subunit bound ATP in the P1a-state, does the trans-D612/E614 loop of the S5-subunit approach sufficiently to allow triggering of ATP hydrolysis (top right panel). In seam subunits, the trans-D612/E614 loop has fully withdrawn (lower right panels).

**Figure S9.**
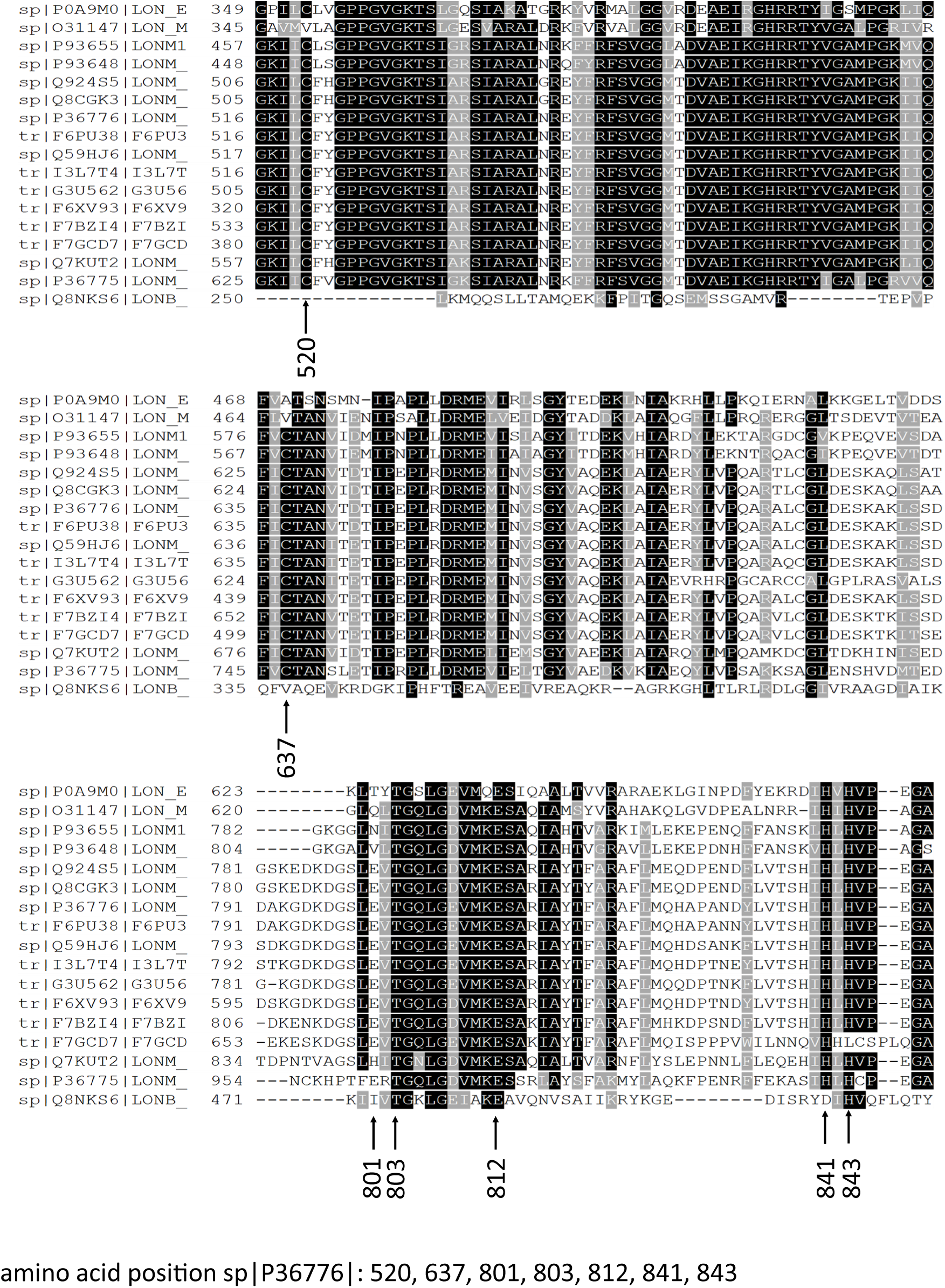
Sequence alignment of LonP1 homologs. Residues C520, C637, E801, T803, E812, H841 and H843 are labelled. Conserved amino acids are indicated.

**Figure S10.**
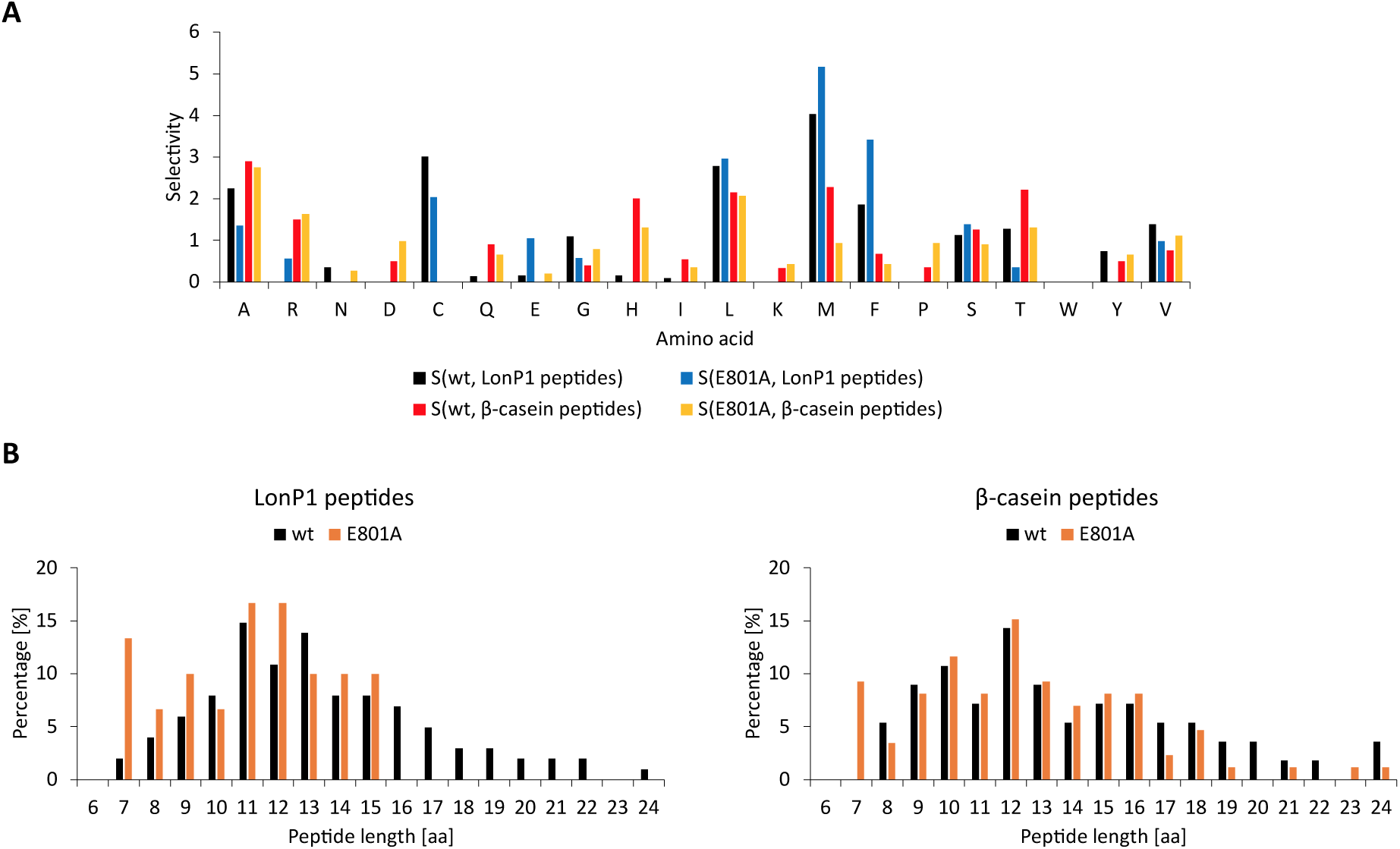
Protease specificity of LonP1 determined by mass spectrometry. (A) Cutting properties (amino acid selectivity S) of LonP1 wt and LonP1 E801A for autocatalysis and FITC-casein (β-casein fraction). The cutting property (selectivity) of LonP1 was calculated as follows, S(aa)=P(aa)/F(aa), whereas P(aa) is the probability of cutting after a given amino acid “aa”. It is calculated as follows: ((number of peptides preceded in sequence by amino acid “aa”) +(number of peptides ending with amino acid “aa”)) / (2 × (total number of peptides)). F(aa) is the frequency of amino acid “aa “occurring in a protein sequence. It is calculated as follows: (number of amino acids “aa “in the protein)/ (total number of amino acids in the protein). (B) Peptide length distribution of LonP1 and β-casein peptides.

**Figure S11.**
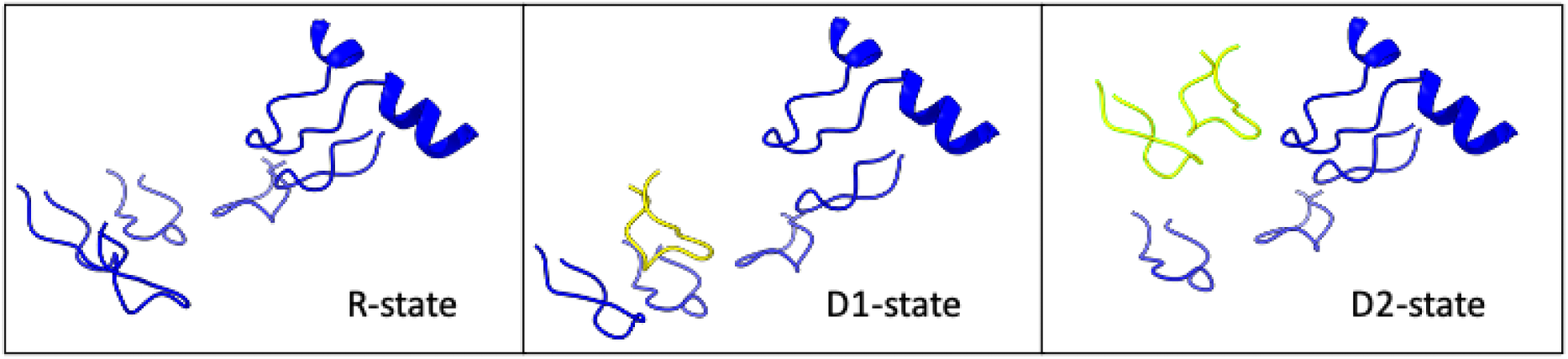
Conversion from the R-state to the P-state through the D1- and D2-states. The R-state subunits were simultaneously fitted as rigid bodies into D1- and D2-state scattering potential maps using the rigid body routines in Phenix-1.18. The resulting assemblies were superimposed using the coordinates of the righthand R-state gap-subunit. The left-hand helical arrangement of the subunits in the R-state is converted into a righthanded arrangement by first lifting up the left-hand gap subunit (in yellow) in the D1-state, then lifting up its left-hand neighbor in the D2-state (both lifted subunits in yellow).

**Figure S12.**
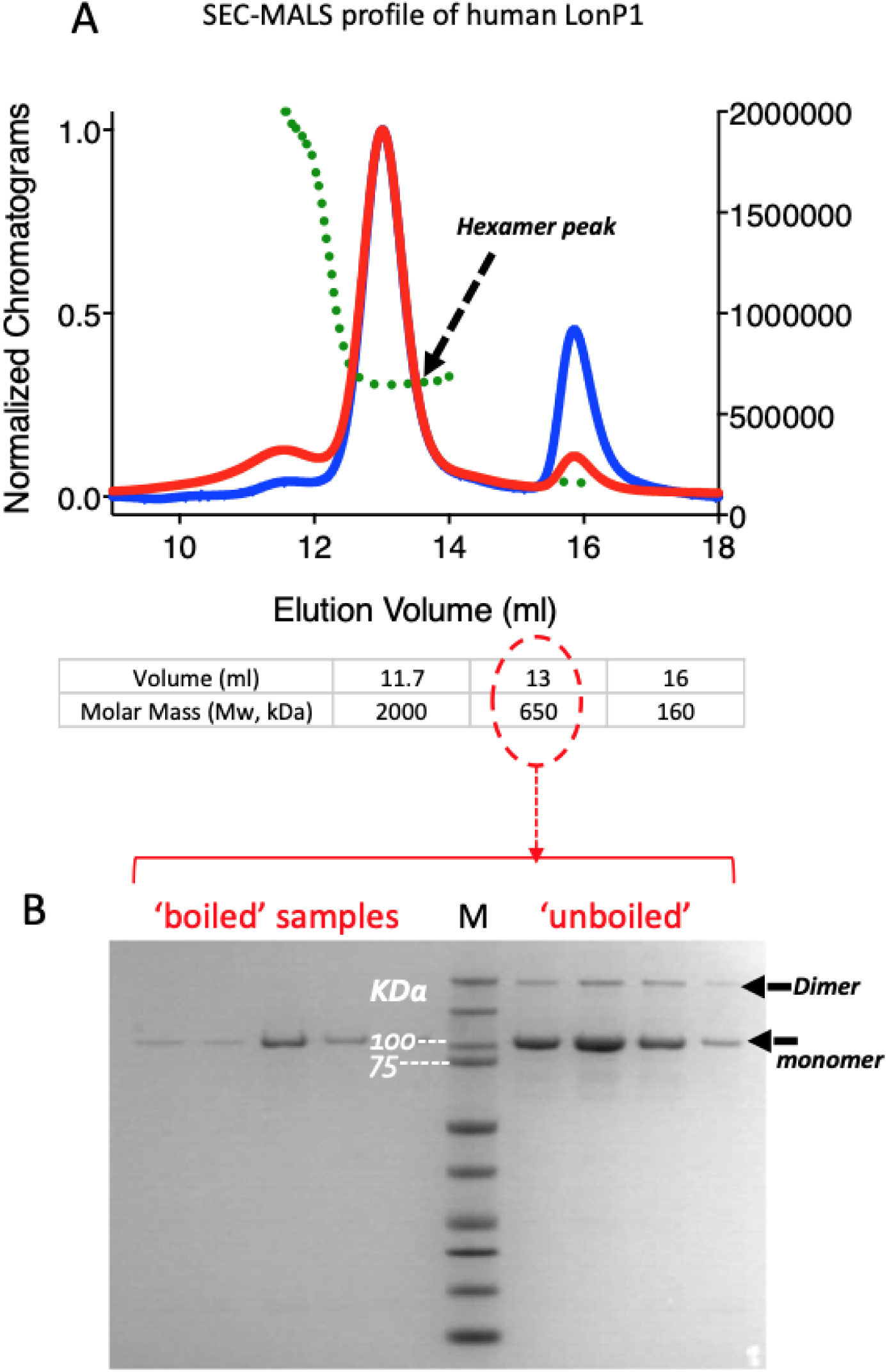
Purification of LonP1. (A) SEC-MALS profile and the SDS-PAGE gel of the peak fraction. The elution volume and the estimated molecular mass of the purified and homogenous peak were 13.0 ml and 650 kDa respectively (from Superose 6 10/30 column, GE healthcare). The bands in (B) shows a single monomer LonP1 near 100 kDa marker and the non-boiled sample indicates a faint band between 200 and 250 kDa.

